# 3D Synaptic Organization of Layer III of the Human Anterior Cingulate and Temporopolar Cortex

**DOI:** 10.1101/2023.02.21.529433

**Authors:** Nicolás Cano-Astorga, Sergio Plaza-Alonso, Javier DeFelipe, Lidia Alonso-Nanclares

**Author notes:** Correspondence to: L. Alonso-Nanclares. **Author contributions** Cano-Astorga: Methodology, Investigation, Formal analysis, Writing - original draft. Plaza-Alonso: Methodology, Formal analysis, Validation, Writing - review & editing. DeFelipe: Conceptualization, Supervision, Funding acquisition, Writing - review & editing. Alonso-Nanclares: Conceptualization, Data curation, Formal analysis, Validation, Supervision, Writing - review & editing. **Competing Interest Statement** The authors declare that they have no competing interest.

## Abstract

The human anterior cingulate and temporopolar cortices have been proposed as highly connected nodes involved in high-order cognitive functions, but their synaptic organization is still basically unknown due to the difficulties involved in studying the human brain. Using Focused Ion Beam/Scanning Electron Microscopy (FIB/SEM) to study the synaptic organization of the human brain obtained with a short post-mortem delay allows excellent results to be obtained. We have used this technology to analyze the neuropil (where the vast majority of synapses are found) of layer III of the anterior cingulate cortex (Brodmann’s area 24) and the temporopolar cortex, including the temporal pole (Brodmann’s area 38 ventral and dorsal) and anterior middle temporal gyrus (Brodmann’s area 21). Our results, based on 6695 synapses fully reconstructed in 3D, revealed that Brodmann’s areas 24, 21 and ventral area 38 showed similar synaptic density and synaptic size, whereas dorsal area 38 displayed the highest synaptic density and the smallest synaptic size. However, the proportion of the different types of synapses (excitatory and inhibitory), the postsynaptic targets and the shapes of excitatory and inhibitory synapses were similar, regardless of the region examined. These observations indicate that certain aspects of the synaptic organization are rather homogeneous, whereas others show specific variations across cortical regions. Since not all data obtained in a given cortical region can be extrapolated to other cortical regions, further studies on the other cortical regions and layers are necessary to better understand the functional organization of the human cerebral cortex.

## Introduction

Numerous studies focusing on the structural organization of the cerebral cortex have been obtained from experimental animals and, in general, it is assumed that these data can be extrapolated to the human cerebral cortex. However, certain fundamental structural and behavioral characteristics are unique to humans and it is therefore imperative to acquire data directly from human brains (e.g., DeFelipe, 2015). Autopsy samples may be the sole source of strictly normal tissue, but the ultrastructure of the post-mortem brain tissue (often obtained 5 or more hours after death) is generally not well preserved, making the tissue unsuitable for detailed quantitative analysis. Therefore, synaptic circuitry data for the normal human brain is virtually non-existent.

However, excellent results have recently been shown in the analysis of human autopsy brain microanatomy after applying a variety of techniques to study brain tissue if it is collected with a post-mortem delay of less than 4 hours. These techniques include intracellular injections to visualize the detailed morphology of neurons (e.g., Benavides-Piccione et al., 2021) and 3D electron microscopy methods using Focused Ion Beam/Scanning Electron Microscopy (FIB/SEM; e.g., Montero-Crespo et al., 2020; Cano-Astorga et al., 2021).

In the present study, we have used FIB/SEM to analyze non-pathological brain tissue samples obtained from autopsy cases with a post-mortem delay of less than 4 h. The goal was to study the normal synaptic organization of the neuropil, where the vast majority of synapses are found (DeFelipe et al., 1999). This technology was chosen because the images obtained are similar to those obtained with transmission electron microscopy, but with the advantage that FIB/SEM permits serial reconstructions of large volumes of tissue to be generated rapidly and automatically (Merchán-Pérez et al., 2009), facilitating detailed 3D reconstructions of synapses and the extraction of synaptic data, even from the human cerebral cortex (Blazquez-Llorca et al., 2013; Domínguez-Álvaro et al., 2018, 2019, 2021a, 2021b; Montero-Crespo et al., 2020, 2021; Cano-Astorga et al., 2021).

We focused on Brodmann’s area (BA) 24, 38 (ventral and dorsal) and 21 (see Zilles and Amunts, 2010), as representative areas of the anterior cingulate (BA24) and temporopolar cortices, including the temporal pole (BA38) and anterior middle temporal gyrus (BA21) (see Mesulam, 2022). It has been proposed that these cortical regions are critically involved in the emergence of a variety of high-order cognitive functions such as memory (Damasio et al., 1990; Wang et al., 2021; Fuster, 2022; Mesulam, 2022); socio-emotional processing and language (Damasio et al., 1996; Kondo et al., 2003; Olson et al., 2013; Xu et al., 2016; Joyce et al., 2022); attention and learning (Morecraft et al., 1993; Keogh et al., 2022); and object/face recognition (Morán et al., 1987; Nakamura and Kubota, 1996; Pascual et al., 2015; Levakov et al., 2021).

Finally, since 3D electron microscopy is very time-consuming, we chose layer III because it plays a key role in the cortico-cortical circuits (Thomson and Lamy, 2007; D’Souza and Burkhalter, 2017), and most of the physiological and morphological studies in the human neurons have been performed in layers II and III (e.g., see Eyal et al., 2018; Gidon et al., 2020; Benavides-Piccione et al., 2021, and references therein). Therefore, the detailed ultrastructural analysis performed in the present study would contribute to a better understanding of this layer of the human cerebral cortex.

## Results

### Volume fraction of cortical elements

Volume fraction (Vv) was estimated applying the Cavalieri principle (Gundersen et al., 1988) in layer III of BA24, vBA38, dBA38 and BA21 to determine the relative volume occupied by different cortical elements: neuropil, cell bodies (including neurons, glial cells and undetermined cells), and blood vessels. The neuropil constituted the main component in all regions (more than 80%; SI Fig. 1; SI Table 1), followed by cell bodies (ranging from 6.80% in vBA38 to 11.88% in BA24) and blood vessels (ranging from 2.81% in BA21 to 3.75% in BA24) (SI Fig. 1; SI Table 1). The volume occupied by cell bodies was significantly higher in BA24 than in any other region (χ^2^, p<0.001; BA24: 11.88%; vBA38: 6.80%; dBA38: 8.09%; BA21: 8.76%).

### Synaptic density

To study the synaptic organization of BA24, vBA38, dBA38 and BA21 across different human cases, their synapses were studied in the 34 stacks of images obtained in the layer III neuropil (i.e., excluding cell bodies, blood vessels, and major dendritic trunks).

A total of 2,052, 1,801, 3,230 and 1,676 synapses were individually identified and 3D reconstructed in BA24, vBA38, dBA38 and BA21, respectively. Of these, 1,430 (BA24), 1,629 (BA38), 2,424 (dBA38), and 1,212 (BA21) synapses were analyzed after discarding incomplete synapses or those touching the exclusion edges of the counting frame (CF; see below) (Table 1; SI Table 2). The synaptic density data were obtained by dividing the total number of synapses included within the CF by its inclusion volume. The synaptic density analyses performed in the neuropil of layer IIIA revealed a very narrow range of values for the number of synapses per volume in all cases and regions analyzed: 0.43–0.55 synapses/μm^3^ in BA24, 0.47–0.62 synapses/μm^3^ in vBA38, 0.65–0.89 synapses/μm^3^ in dBA38, and 0.44–0.51 synapses/μm^3^ in BA21 (Table 1; SI Table 2). The mean synaptic density values obtained in dBA38 were significantly higher (ANOVA; P<0.05) than those obtained in BA24, vBA38 and BA21 (Fig. 1).

**Table 1.** Accumulated data obtained from the ultrastructural analysis of neuropil from layer III of BA24, vBA38, dBA38 and BA21 in human autopsy samples. Data in parentheses have not been corrected for shrinkage. The data for individual cases are shown in SI Table 2. AS: asymmetric synapses; BA: Brodmann’s area; CF: counting frame; d: dorsal; SAS: synaptic apposition surface; SD: standard deviation; SE: standard error of the mean; SS: symmetric synapses; v: ventral.

|  | **BA24** | **vBA38** | **dBA38** | **BA21** |
| --- | --- | --- | --- | --- |
| No. AS | 1335 | 1544 | 2279 | 1129 |
| No. SS | 95 | 85 | 145 | 83 |
| No. synapses<br>(AS+SS) | 1430 | 1629 | 2424 | 1212 |
| % AS | 93.21% | 94.65% | 93.95% | 93.28% |
| % SS | 6.79% | 5.35% | 6.05% | 6.72% |
| CF volume<br>($\mu\text{m}^3$) | 3,144<br>(2,831) | 3,110<br>(2,801) | 3,166<br>(2,851) | 2,483<br>(2,236) |
| No. AS/$\mu\text{m}^3$<br>(mean ± SD) | 0.46±0.06<br>(0.51±0.06) | 0.49±0.09<br>(0.55±0.10) | 0.71±0.11<br>(0.79±0.13) | 0.46±0.06<br>(0.51±0.07) |
| No. SS/$\mu\text{m}^3$<br>(mean ± SD) | 0.03±0.01<br>(0.03±0.01) | 0.03±0.00<br>(0.03±0.00) | 0.05±0.01<br>(0.05±0.01) | 0.03±0.01<br>(0.03±0.01) |
| No. all<br>synapses/$\mu\text{m}^3$<br>(mean ± SD) | 0.49±0.06<br>(0.54±0.06) | 0.52±0.08<br>(0.58±0.09) | 0.76±0.12<br>(0.84±0.13) | 0.49±0.06<br>(0.54±0.07) |
| Intersynaptic<br>distance<br>(nm; mean ± SD) | 845±27<br>(785±25) | 821±28<br>(763±26) | 739±48<br>(686±44) | 821±52<br>(797±50) |
| Area of SAS AS<br>(nm<sup>2</sup>; mean ± SE) | 119,498± 2,425<br>(111,133±2,255) | 111,388±2,298<br>(103,590±2,137) | 77,265±1,510<br>(71,856±1,404) | 113,178±2,626<br>(105,256±2,442) |
| Area of SAS SS<br>(nm<sup>2</sup>; mean ± SE) | 67,537±5,622<br>(62,890±5,229) | 55,084±3,892<br>(51,228±3,620) | 52,460±2,688<br>(48,788±2,500) | 62,469±5,342<br>(58,096±4,968) |

**Figure 1.**
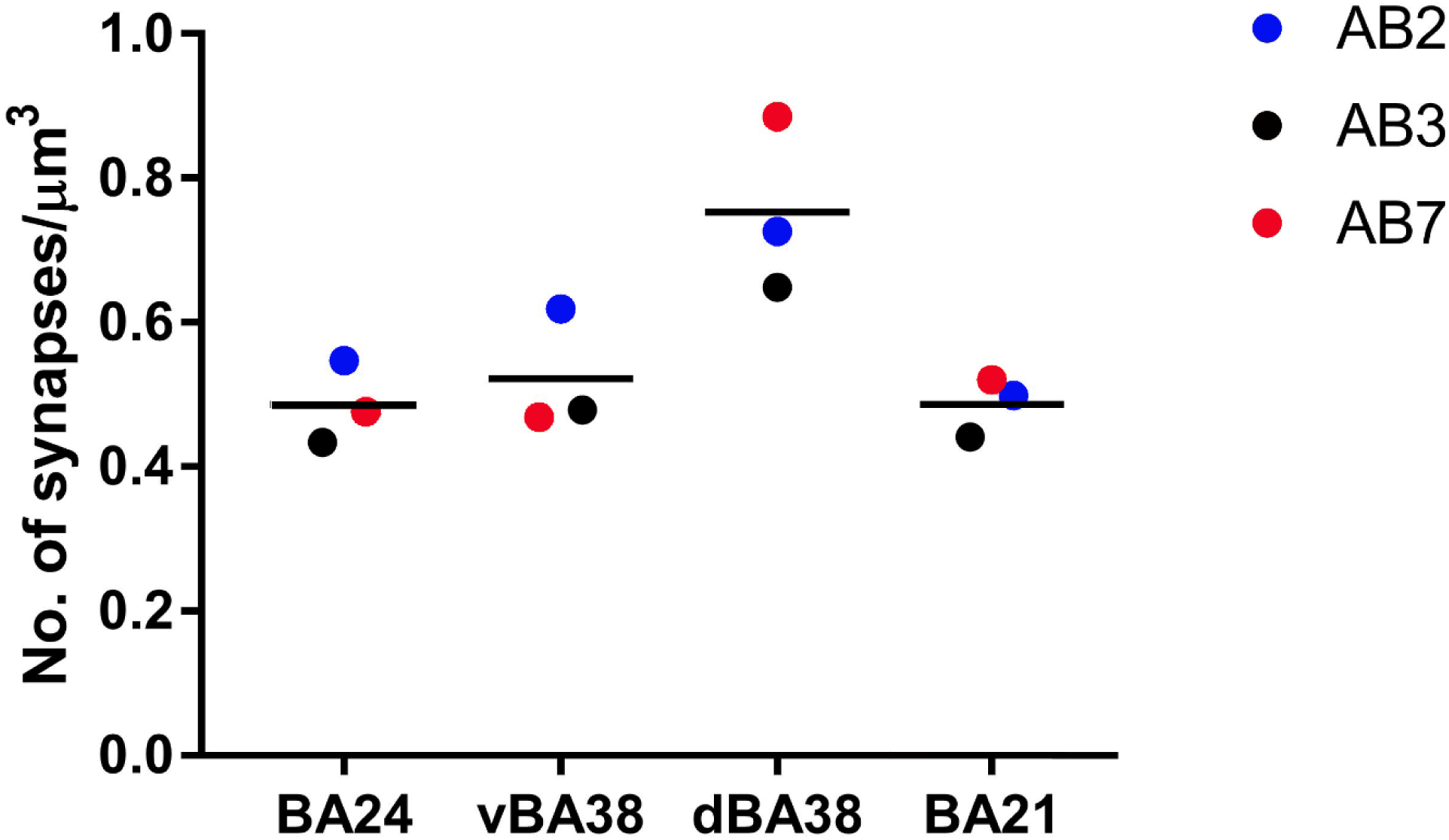
Mean synaptic density of the neuropil from layer III of BA21, BA24, vBA38 and dBA38. Each dot represents a single autopsy case according to the colored key on the right. dBA38 shows a significantly higher synaptic density than the other analyzed regions (ANOVA; P <0.05). BA: Brodmann’s area; d: dorsal; v: ventral.

The proportions of AS and SS were also calculated for all samples. As shown in Table 1 (see also SI Table 2), the proportion of AS:SS was 93:7 in BA24 and BA21, 95:5 in vBA38, and 94:6 in dBA38. No differences were observed in the AS:SS proportion between regions (χ^2^; P>0.05).

### Three-dimensional spatial synaptic distribution

To analyze the spatial distribution of the synapses, the actual position of each of the synapses in each stack of images was compared with the Complete Spatial Randomness (CSR) model. For this, the functions G, K, and F were calculated in the 34 stacks of images analyzed in BA24, vBA38, dBA38 and BA21. We found that 6 stacks of images did not fit into the CSR model, showing a slight tendency for a regular pattern in at least one of the functions (SI Fig. 2A). In the remaining samples (i.e., 28 out of 34 stacks), the 3 spatial statistical functions resembled the theoretical curve that simulates the random spatial distribution pattern, which indicated that synapses fitted a random spatial distribution model in all areas examined (BA24, BA38v, dBA38d and BA21) (SI Fig. 2B).

In addition, the mean distance of each synapse centroid to its nearest neighboring synapse was estimated. The analysis showed that the distance to the nearest neighboring synapse in dBA38 was smaller than in the other cortical areas — differences which were statistically different (KW; P<0.01) compared to BA24 (Table 1; SI Table 2). No significant differences were found between the remaining cortical areas.

### Study of the morphology of the synapses

#### Synaptic size

The study of the synaptic size was carried out analyzing the area of the synaptic apposition surface (SAS) of each synapse identified and 3D reconstructed in all FIB/SEM stacks. As shown in Fig. 2A, B, the size and shape of the SAS were rather variable. The average size of the synapses (measured by the area of the SAS) showed that AS were statistically larger than SS in all regions (Table 1; SI Table 2). To characterize the data distribution of AS and SS SAS area, we performed goodness-of-fit tests to find the theoretical probability density functions that best fitted the empirical distributions of SAS areas in all regions. We found that the best fit corresponded to log-normal distributions, with some variations in the location (µ; range: 10.888 – 11.413 in AS and 10.673 – 10.820 in SS) and scale (σ; range: 0.7872 – 0.8690 in AS and 0.6499 – 0.8091 in SS) parameters). This was the case in all regions for both AS and SS, although the fit was better for AS than for SS, probably due to the smaller number of SS.

**Figure 2.**
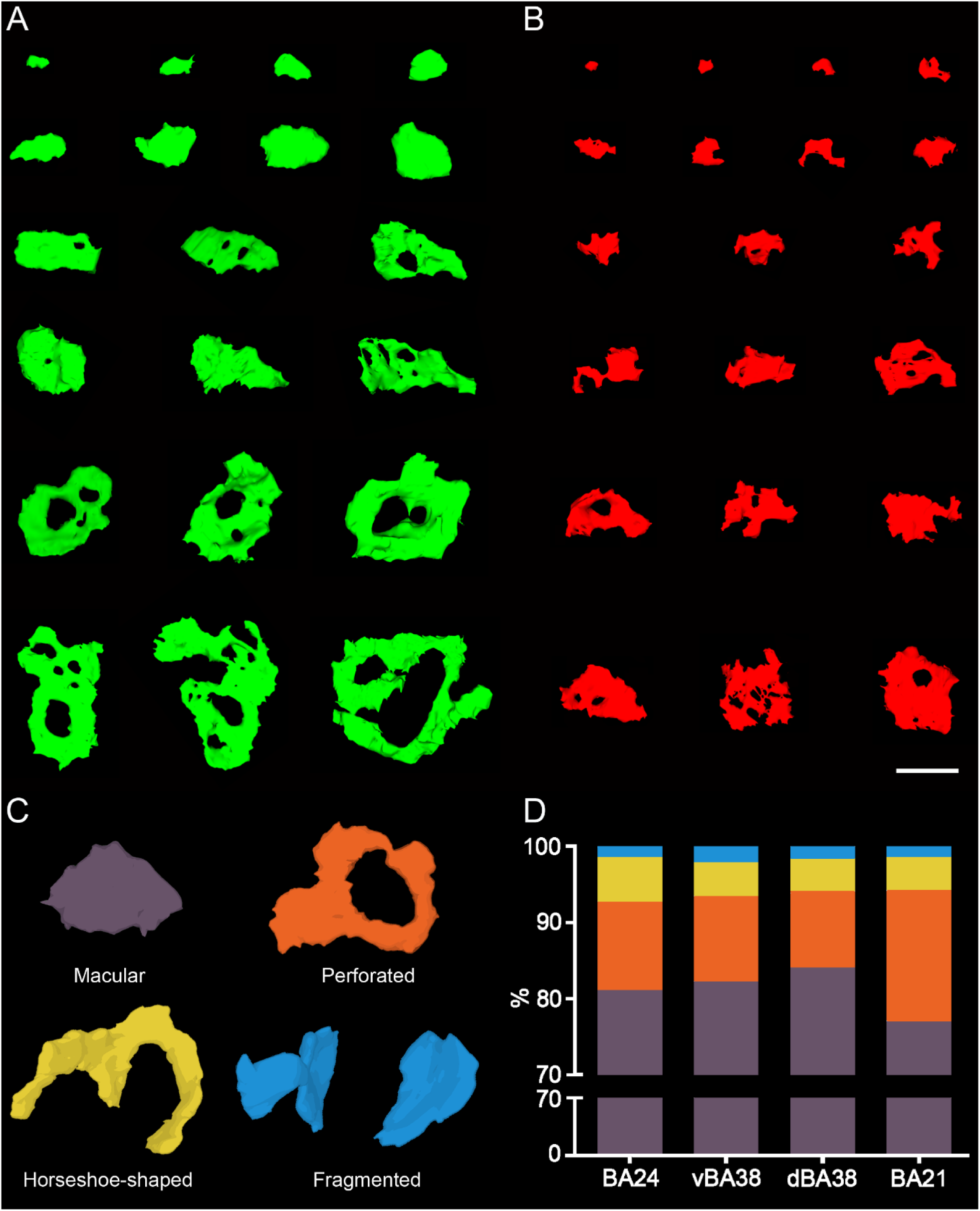
3D Synaptic morphology representation. (A, B) Representative examples of SAS of asymmetric synapses (A, green) and symmetric synapses (B, red). Analyses of SAS were distributed into 20 bins of equal size (an example of each bin has been illustrated). (C) Schematic representation of the shape of the synaptic junctions: macular synapses, with continuous disk-shaped PSD; perforated synapses, with holes in the PSD; horseshoe-shaped synapses, with tortuous horseshoe-shaped perimeter with an indentation; and fragmented synapses, with two or more PSDs with no connections between them. (D) Proportion of macular, perforated, horseshoe-shaped, and fragmented AS in BA24, vBA38, dBA38 and BA21. BA21 showed a higher proportion of complex-shaped synapses (including perforated, horseshoe, and fragmented synapses) than BA24, vBA38 and dBA38 (χ^2^; P<0.0001). Scale bar (in B) indicates 500 nm in A and B. BA: Brodmann’s area; d: dorsal; SAS: synaptic apposition surface; v: ventral.

Comparison of the probability density functions between AS and SS showed that larger synapses were more frequent in AS than SS in all regions (KS; P<0.001; SI Fig. 3). Upon the analyses of the average area of the SAS of AS between cortical regions, significant differences (KW; p<0.0001) were found between regions, indicating that AS were smaller in dBA38 than in vBA38, BA24 and BA21. Also, significant differences (KW; p<0.05) were found between the mean SAS area of AS in vBA38 and BA24, indicating that AS in vBA38 were smaller than in BA24 (Table 1; SI Table 2). Similarly, significant differences were found when comparing the AS SAS area frequency distribution between cortical regions, indicating that smaller AS were more frequent in dBA38 than vBA38, BA24 and BA21 (KS, P<0.0001). Furthermore, smaller AS were more frequent in vBA38 than BA24 (KS, P=0.0004). The number of SS examined in all areas was not sufficient to perform a robust statistical analysis focusing on possible differences between SAS in different regions.

#### Synaptic shape

To analyze the shape of the synaptic contacts, we classified each identified synapse into four categories: macular (with a flat, disk-shaped postsynaptic density (PSD)), perforated (with one or more holes in the PSD), horseshoe (with an indentation in the perimeter of the PSD), or fragmented (with two or more physically discontinuous PSDs) (Fig. 2C; for a detailed description, see Santuy et al., 2018a; Domínguez-Álvaro et al., 2019). The analyses showed that the vast majority of AS presented macular shape in all of the cortical regions analyzed (range: 77–84%), followed by perforated (range: 10–17%), horseshoe (range: 4–6%) and fragmented (range: 1–2%; SI Table 3). Regarding the SS, the results were very similar in all analyzed regions: the majority of SS presented a macular shape (range: 85–92%; SI Table 3), whereas the perforated (range: 2–8%) and horseshoe (range: 2–6%) shapes were less frequent. SS exhibiting a fragmented shape were scarce; indeed, this shape was only identified in 3 synapses in vBA38, 1 synapse in dBA38 and 1 synapse in BA21 (SI Table 3).

To determine whether there was a difference in the proportion of AS shapes between regions, contingency tests were applied. Perforated, horseshoe, and fragmented synapses were computed as a whole (as complex-shaped synapses), and statistically significant differences were found (χ^2^, P<0.0001) indicating that complex-shaped AS were more frequent in BA21 (33.0%) than in vBA38 (17.8%), dBA38 (15.9%) and BA24 (18.8%) (Fig.2D; SI Table 3). Concerning SS, the number of synapses was not sufficient to perform a robust statistical analysis to analyze possible differences between cortical areas.

#### Synaptic size and shape

We also determined whether the shape of a synapse is related to its size. For this purpose, the areas of the SAS —AS and SS— were analyzed according to the synaptic shape. We found that the area of the macular AS was smaller (KW; P<0.0001) than the area of the perforated, horseshoe, and fragmented AS in all regions (SI Table 4;). Similarly, the smaller AS were more frequent in the macular-shaped synapses in all cortical regions (KS; P<0.0001; SI Fig. 4). SS numbers were not sufficient to perform a robust statistical analysis.

### Study of the postsynaptic targets

Postsynaptic targets were identified and classified as dendritic spines (axospinous synapses) or dendritic shafts (axodendritic synapses). We also determined whether the synapse was located on the neck or head of a spine (SI Fig. 5).

When the postsynaptic element was identified as a dendritic shaft, it was classified as “with spines” or “without spines” (SI Table 5). The postsynaptic elements of 1,262, 1,574, 2,350 and 1,143 synapses were determined in BA24, vBA38, dBA38 and BA21, respectively. In all analyzed cortical regions, most synapses were AS established on dendritic spine heads (range: 68.5–73.4%) followed by AS on dendritic shafts (range: 20.8–24.0%), SS on dendritic shafts (range: 3.7–5.7%) and AS on dendritic spine necks (range: 0.6–1.1%) (Fig. 3). The least frequent types of synapses were SS established on dendritic spine heads (range: 1.1–1.3%) and SS on dendritic spine necks (range: 0.3–0.6%) (Fig. 3).

**Figure 3.**
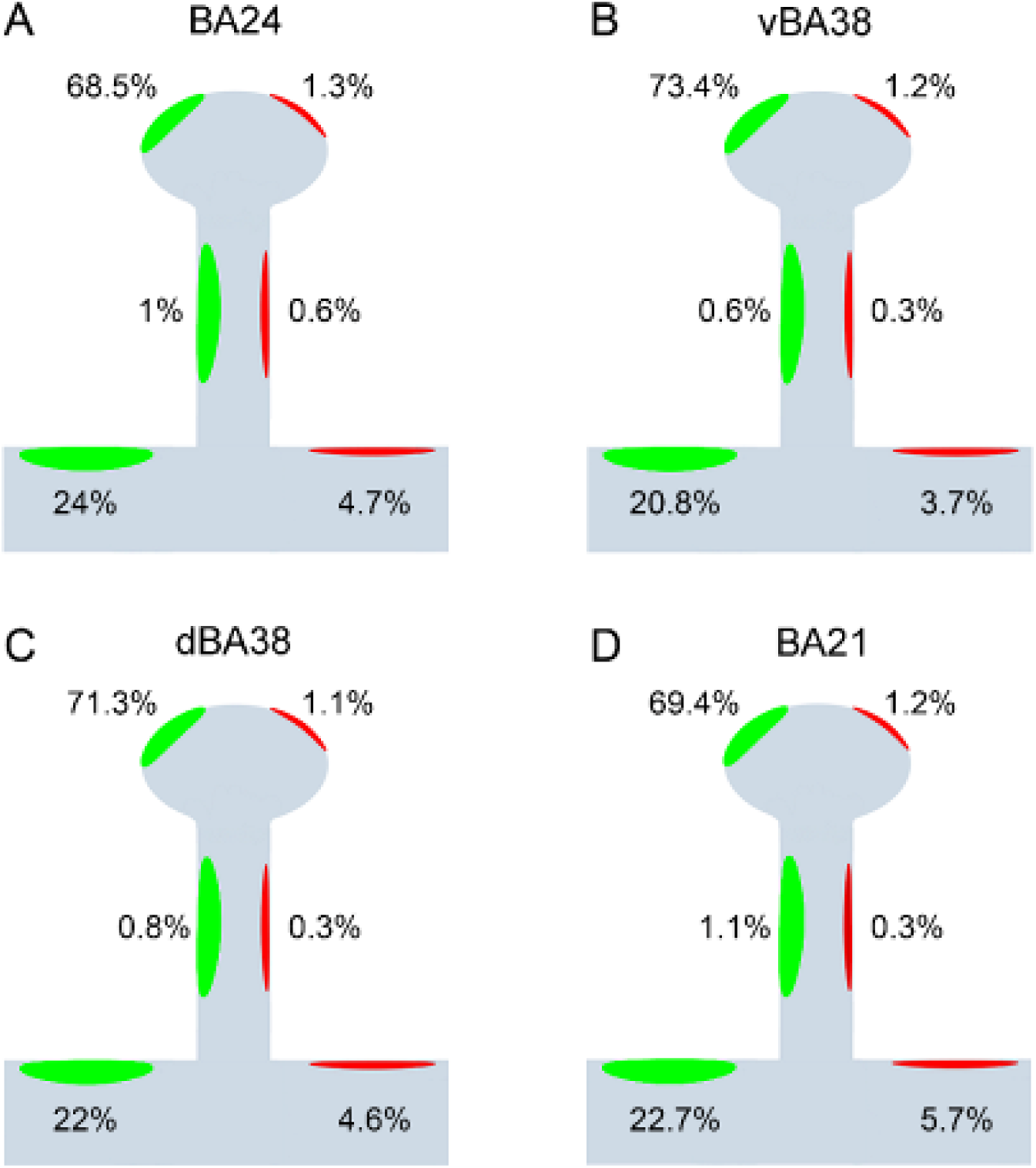
Schematic representation of the asymmetric and symmetric synapses on the different postsynaptic targets (spine head, spine neck and dendritic shaft) from analyzed synapses in layer III of BA24 (A), vBA38 (B), dBA38 (C) and BA21 (D). Spine head percentages include both complete and incomplete spines. Dendritic shaft percentages include dendritic shafts with and without spines. No significant differences (χ^2^; P>0.05) were found between cortical regions. AS have been represented in green and SS in red. BA: Brodmann’s area; d: dorsal; v: ventral.

Comparisons of the AS postsynaptic targets between cortical regions were analyzed and no differences were found (χ^2^, P>0.05; SI Table 5). Concerning SS, the number of synapses examined was not sufficient to perform a robust statistical analysis.

In addition, to detect the presence of multiple synapses, an analysis of the spine heads was performed to determine the number and type of synapses established on them. The vast majority of spines had a single AS (range: 93-96%) followed by spines with a large variety in the synapse number and location (head or neck) of AS and SS (Fig. 4). A comparison of the multisynaptic spine proportions between cortical regions was also carried out. Spines with more than one AS (Fig. 4) were significantly more frequent in BA24 than in the other regions (χ^2^, P<0.05).

**Figure 4.**
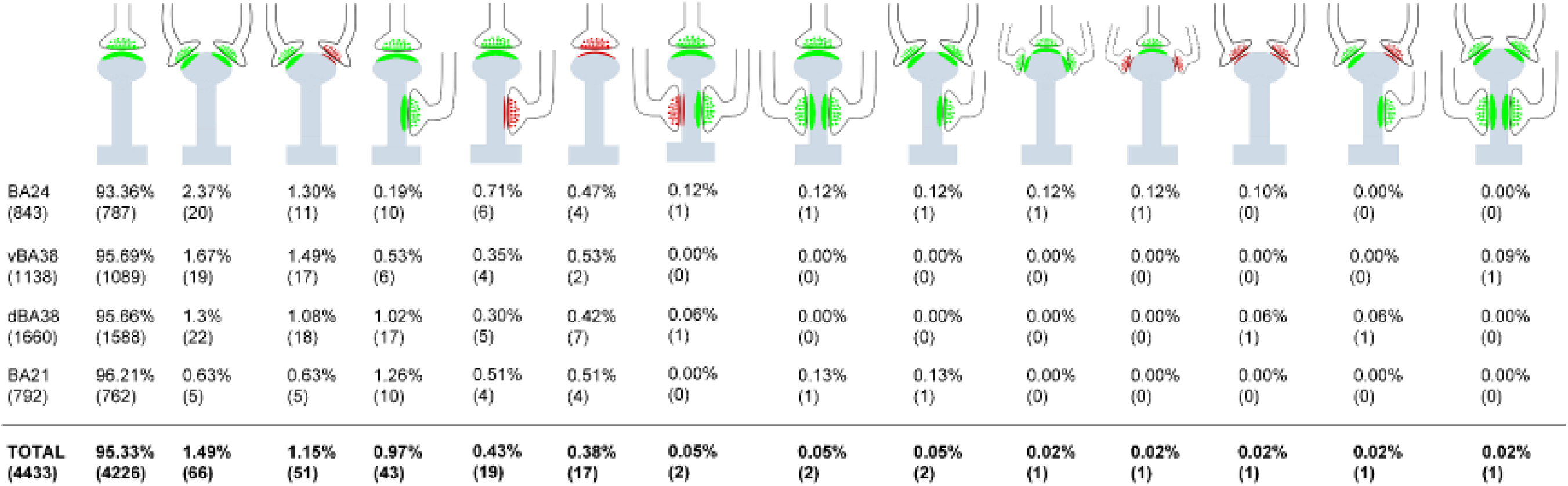
Schematic representation of dendritic spine heads (including both complete and incomplete spines) receiving single and multiple synapses in BA24, vBA38, dBA38 and BA21. Percentages of each type are indicated (absolute numbers of synapses are in parentheses). AS have been represented in green and SS in red. Spines with more than one AS (including all combinations) were more frequent in BA24 than in the other regions (χ^2^, P<0.05). AS: asymmetric synapses; BA: Brodmann’s area; d: dorsal; SS: symmetric synapses; v: ventral.

#### Postsynaptic targets and synaptic size

Finally, we also determined whether the postsynaptic elements of the synapses were related to their size. For this purpose, the areas of the SAS —of both AS and SS— were analyzed according to the postsynaptic targets. In order to perform a more accurate analysis of the size of the axospinous synapses, we excluded those synapses established on incomplete spines. Large AS were more frequently found on spines than on dendritic shafts (with and without spines) in all regions (KS; P<0.01; SI Fig. 6).

The area of the AS established on spines was significantly larger than the area of the AS established on different targets depending on the region (KW; P<0.01; SI Table 6). The number of SS analyzed was not sufficient to perform a robust statistical analysis.

### Synaptic interindividual variability

We also studied the variability between individuals. As shown in SI Table 2, the synaptic density found in the three cases in dBA38 was more variable (0.65–0.89 synapses/µm^3^) than in any other cortical region, especially when comparing to BA21, where the synaptic density of the three cases were remarkably similar (0.44–0.51 synapses/µm^3^). In the case of dBA38, the percentage of macular synapses was between 83 and 94%, which is higher than any other cortical region and, again, notably higher than in BA21 where the macular synapse percentages were very similar (75.9–77.6%; SI Table 3) in the three cases. Finally, in BA21 the percentage of AS established on dendritic spine heads varied depending on the case: 67%, 71% and 82% in cases AB3, AB2 and AB7, respectively. However, in BA24, the percentages for the three cases were very similar (71–74%; SI Table 5).

## Discussion

The present results provide a large quantitative ultrastructural dataset of synapses in layer III of the anterior cingular and temporopolar cortex using 3D EM. The main result is that there were synaptic characteristics specific to particular regions (synaptic density and synaptic size were different in dBA38), whereas some other synaptic characteristics were common to all analyzed regions, including: i) a similar AS:SS ratio; ii) synapses fitted into a random spatial distribution; iii) the SAS area of AS was larger than that of SS; iv) most synapses displayed a macular shape, and they were smaller than complex-shaped synapses; v) most synapses were AS established on dendritic spines, followed by AS on dendritic shafts and SS on dendritic shafts; vi) AS on spine heads were larger than AS on dendritic shafts; and vii) most dendritic spine heads receive a single AS.

### Synaptic Density

Most connections in the cerebral cortex are established by point-to-point chemical synapses (DeFelipe, 2015). Therefore, determining the synaptic density in a given region is important since this parameter is crucial in terms of connectivity and functionality. In the present study, we found that the mean synaptic density of the neuropil of layer III in dBA38 (0.79 synapses/μm^3^) was significantly higher than in BA24 (0.49 synapses/μm^3^), BA21 (0.49 synapses/μm^3^) and vBA38 (0.52 synapses/μm^3^). Previous studies using the same techniques (in the same autopsy cases, AB2 and AB3) have shown that the synaptic density in the neuropil of other cortical regions —such as layer II of the transentorhinal cortex (Domínguez-Álvaro et al., 2018) and layers II and III of the entorhinal cortex (Domínguez-Álvaro et al., 2021a)— were similar to the synaptic density of BA24, BA21 and vBA38 found in the present study. However, in the deep and superficial stratum pyramidale of the C*ornu Ammonis* 1 (CA1) field of the hippocampus of these autopsy cases, the synaptic density was 0.67 and 0.99 synapses/μm^3^, respectively (Montero-Crespo et al., 2020), which is closer to the synaptic density found in dBA38. Since the tissue processing and analysis methods were identical, similarities and differences would appear to be attributable to specific characteristics of the brain region and layer analyzed.

The proportion of tissue occupied by neuropil (Vv-neuropil) is clearly also an important factor in terms of synaptic connectivity of the cortical region and layer. In the present study, the Vv-neuropil constituted the main component of layer III of BA24, vBA38, dBA38 and BA21 (range: 84–90%). These were similar values to those obtained in the same autopsy cases in layer II of the human transentorhinal cortex (Domínguez-Álvaro et al., 2018) and in the pyramidal cell layer of the CA1 field of the hippocampus (Montero-Crespo et al., 2020). These apparently small differences in the Vv-neuropil, however, could imply substantial differences in the overall number of synapses (DeFelipe et al., 1999). Since in the present study we found that the synaptic density varies within a range of between approximately 0.5 and 0.8 synapses per µm^3^, both the density and the Vv-neuropil should be considered together. For example, BA24 had a lower Vv-neuropil (84%) than BA21 (88%), dBA38 (89%) and vBA38 (90%). However, the synaptic density of BA24 was similar to BA21 and vBA38, but much lower than in dBA38. Hence, considering the synaptic density and the Vv-neuropil together, layer III of BA24 must contain a lower number of synapses than the other regions. Thus, the significance of volume fraction variations must be considered together with the synaptic density and the total volume of the layer to better understand the overall connectivity of these cortical regions.

Moreover, the present results regarding synaptic density are in line with the higher spine density and pyramidal complexity reported in layer III from BA38 (Benavides-Piccione et al., 2013, 2021). These studies focusing on human pyramidal neurons have shown that in the BA38, pyramidal neurons had higher branching complexity and higher dendritic spine density than in BA21 and BA24 (Benavides-Piccione et al., 2021). In addition, both BA38 and BA21 had higher spine densities than BA24 (Benavides-Piccione et al., 2013). Since pyramidal neurons represent the majority of cortical neurons and their dendritic spines are the major target of AS, a direct correlation between synaptic density and complexity of pyramidal neurons may exist. Further studies would be needed to investigate the possible correlation of dendritic spine density and synaptic density in other cortical areas.

### Proportion of Synapses and Spatial Synaptic Distribution

It is well established that the cortical neuropil has a larger proportion of AS than SS regardless of the cortical region and species (rodent, monkey or human). Using transmission electron microscopy, this proportion varies between 80 and 95% for AS and 20–5% for SS (reviewed in DeFelipe et al., 2002; DeFelipe 2011; for a recent study see Cano-Astorga et al., 2021 and references therein). Similar proportions have been found using FIB/SEM in the present study and in other cortical regions (Domínguez-Álvaro et al., 2018, 2021a; Montero-Crespo et al., 2020; Cano-Astorga et al., 2021). Therefore, regarding the proportion of AS and SS in the human cerebral cortex, it seems that the values are among the highest for AS and among the lowest for SS in the mammalian cerebral cortex.

The significance of this similar AS:SS ratio is difficult to interpret due to the differences in the cytoarchitecture, connectivity, and functional characteristics of different brain regions. A variety of neurochemical and functional types of pyramidal and GABAergic neurons are present in these regions but their densities vary. For example, the human cingulate cortex (BA24) contains more immunoreactive interneurons for tyrosine-hydroxylase than the temporal cortex (including BA21 and BA38; Benavides-Piccione et al., 2005; Benavides-Piccione and DeFelipe, 2007), whereas the human temporal pole cortex (BA38) has more immunoreactive interneurons for parvoalbumin than BA21 and BA24 (Blázquez-Llorca et al., 2010).

One possibility is that the AS:SS ratio in the dendritic arbor of the different types of neurons may be similar. However, in other cortical regions of different species, it has been shown that there are differences in the number of GABAergic and glutamatergic inputs in several neuronal types (e.g., DeFelipe and Fariñas 1992; Freund and Buzsáki 1996; DeFelipe 1997; Somogyi et al., 1998; Schubert et al., 2007; Markram et al., 2015; Tremblay et al., 2016; Hu et al., 2018; Sohal and Rubenstein, 2019; Chini et al., 2022). Thus, it would be necessary to examine the synaptic inputs of each particular cell type to determine differences in their AS:SS ratio, even though the overall AS:SS proportion does not vary in the neuropil.

The analysis of the spatial organization of synapses showed that the synapses were randomly distributed in the neuropil from most of the samples of BA24, vBA38, dBA38 and BA21. This spatial distribution has also been found in other regions of the human brain, including transentorhinal and entorhinal cortex, and CA1 field of the hippocampus (Domínguez-Álvaro et al., 2018, 2021a; Montero-Crespo et al., 2020). Therefore, the present results —regarding the AS:SS proportion and spatial distribution of synapses— support the idea that the synaptic organization of the human cerebral cortex follows general rules.

### Synaptic Size and Shape

It has been proposed that synaptic size correlates with release probability, synaptic strength, efficacy, and plasticity (see Chindemi et al., 2022 and references therein). Several methods have traditionally been used to estimate the size of synaptic junctions making it difficult to compare between different studies (reviewed in Cano-Astorga et al., 2021). In addition, most studies estimate the size of synapses in general (mostly AS) but few reports provide specific data regarding the size of the SS. Here, we have found that AS were larger than SS in all the regions analyzed, as previously reported in other human cortical regions such as layer II of the transentorhinal cortex (Domínguez-Álvaro et al., 2018), layers II and III of the entorhinal cortex (Domínguez-Álvaro et al., 2021a), and in all layers of the CA1 field of the hippocampus (Montero-Crespo et al., 2020). Using the same techniques in other mammalian species, AS were also found to be larger than SS in all layers of the somatosensory cortex of the Etruscan Shrew (Alonso-Nanclares et al., 2022) and juvenile rat (Santuy et al., 2018a). However, it has been shown that SS were larger than AS in certain layers of the CA1 field of the mouse hippocampus and in certain layers of the mouse somatosensory cortex (Santuy et al., 2020; Turégano-López, 2022). Thus, further studies should be performed in other cortical areas, layers and species to find out if this characteristic is a regional and/or species-specific rule of the mammalian cerebral cortex.

In addition, the densities of N-methyl-D-aspartate (NMDA), kainate, and norepinephrine receptors have been reported to be higher in BA24 than in BA38 (Palomero-Gallagher et al., 2008; Zilles and Palomero-Gallagher, 2017). This seems to be in contrast with our results since synaptic density in BA24 was lower than in dBA38. Synaptic size is related to the number of receptors in the PSD — larger PSD have a larger number of receptors (reviewed in Lüscher et al., 2000; Lüscher and Malenka 2012; Toni et al., 2001; Magee and Grienberger, 2020; Sumi and Harada, 2020). In particular, BA24 had the largest AS compared to the other regions examined. Nevertheless, the above-mentioned receptors have been reported to be synaptically and extra-synaptically located depending on each receptor (Palomero-Gallagher and Zilles, 2019). Therefore, caution should be taken regarding possible correlations between receptor densities, number of synapses and/or synaptic size. Indeed, the differences and similarities in the synaptic density between human cortical regions —as well as the particular regional densities of receptors of various neurotransmitters— should be considered together to better understand the functional organization of synaptic circuits.

Furthermore, we found that most synapses presented a macular shape (81–85%), whereas 15–19% were complex-shaped synapses (including perforated, horseshoe and fragmented) — with complex-shaped AS being larger than macular AS. These observations are comparable to previous reports in other brain areas and species (Geinisman et al., 1987; Jones et al., 1991; Neuman et al., 2016; Hsu et al., 2017; Calì et al., 2018; Santuy et al., 2018a; Domínguez-Álvaro et al., 2019, 2021a; Montero-Crespo et al., 2020; Cano-Astorga et al., 2021). It has been reported that complex-shaped synapses have more α-amino-3-hydroxy-5-methyl-4-isoxazolepropionic acid (AMPA) and NMDA receptors than macular synapses, and they are thought to constitute a relatively ‘powerful’ population of synapses with more long-lasting memory-related functionality than macular synapses (Geinisman et al., 1987, 1991, 1992a, 1992b, 1993; Lüscher et al., 2000; Toni et al., 2001; Ganeshina et al., 2004a, 2004b; Spruston 2008). Thus, determining the shape of synapses is interesting from the functional point of view. The proportion of complex-shaped synapses was higher in BA21 than in the other analyzed regions. In particular, dBA38 had the lowest proportion of complex-shaped synapses. Thus, dBA38 seems to display a variety of distinct synaptic characteristics compared to vBA38, BA21 and BA24, including high synaptic density, small synaptic size (SAS area) and a low proportion of complex-shaped synapses.

### Postsynaptic Targets

A clear preference of glutamatergic axons (forming AS) for spines, and GABAergic axons (forming SS) for dendritic shafts, was observed (Fig. 3). This is in line with the numerous studies using electron microscopy in a variety of cortical regions and species (reviewed in DeFelipe et al., 2002). However, this characteristic is commonly misinterpreted as implying that synapses on shafts are mostly SS. In fact, this is not the case — quantitative analyses of synapses in the neuropil have shown that most synapses are AS established on dendritic spines, followed by AS on dendritic shafts, and then SS on dendritic shafts (Beaulieu et al., 1992; Peters et al., 2008; Hsu et al., 2017; Calì et al., 2018; Santuy et al., 2018b; Yakoubi et al., 2019; Montero-Crespo et al., 2020, 2021; Domínguez-Álvaro et al., 2019, 2021a, 2021b; Cano-Astorga et al., 2021).

In the present study, the simultaneous analysis of the synaptic type and postsynaptic target showed that the proportions of AS on spines (“axospinous”) were around 73–77%. Using the same FIB/SEM technology in other cortical regions in the same autopsy samples (AB2 and AB3), the proportions of AS established on spines were similar in layer II of the human transentorhinal cortex (75%; Domínguez-Álvaro et al., 2021a) and stratum oriens, deep stratum pyramidale, and stratum radiatum of the CA1 hippocampal field (77–81%; Montero-Crespo et al., 2020). However, this percentage was much lower in layers II and III of the human entorhinal cortex (60% and 56%, respectively; Domínguez-Álvaro et al., 2021a) and in the stratum lacunosum-moleculare of the CA1 hippocampal field (57%; Montero-Crespo et al., 2020). The superficial stratum pyramidale of the CA1 hippocampal field had a higher proportion of AS established on spines (88%; Montero Crespo et al., 2020). Therefore, these peculiarities in the proportion of AS on spines represent another microanatomical specialization of the cortical regions (and layers) examined which may have important functional implications in the processing of information of these cortical regions.

In addition, we found that AS established on dendritic spine heads were larger than those established on dendritic shafts or dendritic spine necks, as was previously reported in layer II of the transentorhinal cortex (Domínguez-Álvaro et al., 2019) and layer II and III of the entorhinal cortex (Domínguez-Álvaro et al., 2021a). However, in the CA1 field of the hippocampus, no differences were found between the axospinous and axodendritic synaptic sizes (Montero-Crespo et al., 2020). Again, this suggests another regional synaptic specialization.

Regarding the number of synapses per dendritic spine, most dendritic spines established a single AS (Fig. 4), which is similar to layer II of the transentorhinal cortex (94.5%; Domínguez-Álvaro et al., 2019). However, this percentage was lower in layers II and III of the entorhinal cortex (90.3% and 89.6%, respectively; Domínguez-Álvaro et al., 2021a) and higher in the CA1 field of the hippocampus (97.7%; Montero-Crespo et al., 2020). Thus, these differences and similarities in the proportion of spines establishing single or multiple synapses between cortical regions could indicate another specialization feature of the human cerebral cortex.

The functional relevance of the presence of single/multiple synapses on the same spine remains to be elucidated. In the mice neocortex, it has been proposed that spines receiving one AS and one SS are electrically more stable than spines establishing a single AS (Villa et al., 2016). Furthermore, it has been proposed that SS may play a key role in spine calcium signaling (Chiu et al., 2013 and Kleinjan et al., 2023). Since the vast majority of spines are single innervated spines, it is possible that multisynaptic spines represent a special postsynaptic target. For example, in the monkey and human cerebral cortex, it has been shown that a particular type of GABAergic cell (forming SS), called double bouquet cells, made up around 38-48% of synapses on spines that receive additional AS (Somogyi and Cowey, 1981; DeFelipe et al., 1989, 1990; De Lima and Morrison, 1989; del Rio and DeFelipe, 1995; Lukacs et al., 2022). This is a remarkable high percentage of SS made on spines considering that dual innervated spines represent only 1–2% of the total population of synapses found in the neuropil.

### Synaptic organization and connectivity

Since layer III is particularly relevant regarding cortico-cortical circuits (Felleman and Van Essen, 1991; Thomson and Lamy, 2007; Barbas, 2015; D’Souza and Burkhalter, 2017; Rockland, 2019), the differences in synaptic characteristics between BA24, vBA38, dBA38 and BA21 may be related to the diversity of cortical circuits in these regions (SI Fig. 7). For example, it has been described that the posterior cingulate cortex and the dorsolateral prefrontal cortex constitute a major input to BA24, vBA38 and BA21, but not to dBA38 (Pascual et al., 2015; Jackson et al., 2016; Dezachyo et al., 2021). In addition, vBA38 and dBA38 share inputs that are absent in BA24 and BA21, such as those coming from the orbital prefrontal cortex, ventrolateral prefrontal cortex and the supramarginal gyrus. BA24 and BA21, on the other hand, receive inputs from the insular cortex, which are absent in vBA38 and dBA38 (Pascual et al., 2015; Jackson et al., 2016; Dezachyo et al., 2021). Moreover, each region is characterized by the selectivity of their inputs: BA24 preferably receives inputs from the primary motor cortex and the parietal operculum cortex; vBA38 preferably receives inputs from the medial prefrontal cortex, hippocampal formation, posterior parahipocampal region, entorhinal and perirhinal cortex, and fusiform gyrus; and dBA38 preferably receives inputs from the anterior cingulate cortex, superior temporal gyrus and the neighboring areas of the Heschel’s gyrus and postcentral gyrus (Pascual et al., 2015; Jackson et al., 2016; Dezachyo et al., 2021). In addition, as illustrated in SI Fig. 7, there are differences in the interhemispheric connectivity in the analyzed regions. It has been proposed that there are hemispheric preferences regarding the high-order functions of the cerebral cortex, especially in humans (Bartolomeo and Seidel Malkinson, 2019). It has been reported that the connectivity of vBA38 and dBA38 mainly occurs in the ipsilateral hemisphere, whereas BA24 and BA21 showed a more bilateral connectivity pattern (Pascual et al., 2015; Jackson et al., 2016; Dezachyo et al., 2021). Thus, differences found regarding synaptic organization between BA24, vBA38, dBA38 and BA21 may be related to the structural and functional differences of the regions connected with them (for a detailed description, see Zachlod et al., 2022 and references therein).

Furthermore, functional and structural connectivity studies on human brain have proposed that the anterior cingulate (BA24), the temporal pole (BA38) and the middle temporal (BA21) cortices are equally connected nodes (Sporns and Zwi, 2004; Sporns et al., 2005; Van Essen et al., 2013; Paquola et al., 2020). However, our present results show a differential synaptic organization in these regions, especially in terms of the synaptic density and synaptic size (see above) of excitatory synapses (which are by far the most common type of synapses involved in corticocortical connections). Nevertheless, comparing connectivity studies with our ultrastructural data is rather difficult since we have focused on layer III neuropil. That is, our results on the synaptic connectivity are restricted to dendritic shafts and dendritic spines of this particular layer of the selected cortical regions, whereas other cortical layers also receive corticocortical connections (Felleman and Van Essen, 1991; Barbas, 2015; Rockland, 2019). Furthermore, other fundamental aspects of the synaptic circuitry should be considered to decipher the relationship between local synaptic organization and functional and anatomical connectivity between cortical areas. For example, corticocortical axons innervate both excitatory and inhibitory neurons but there are variations in the types and proportions of these neurons. In addition, the perisomatic GABAergic innervation of pyramidal cells (i.e., innervation of the proximal dendrites, soma and axon initial segment) is critical in the control of pyramidal cell action potential output and synchronization. It has been proposed that the inhibitory synapses targeting pyramidal cell somata, dendritic shafts and dendritic spines differs with regard to their synaptic strength (e.g., Miles et al., 1996; Xue et al., 2014; Kubota et al., 2015). In addition, there is a variability in the density and number of perisomatic GABAergic boutons across layers and cortical areas in the human brain (e.g., Inda et al., 2007; Blázquez-Llorca et al., 2010; Ostos et al., 2022). Thus, an integrative multiscale study at micro-, meso-, and macroscopic levels would be necessary to obtain more comprehensive knowledge about the synaptic organization.

Finally, a number of studies highlighted the interindividual variability in the structural and functional organization of the human brain (Jacobs and Scheibel, 1993; Benavides-Piccione et al., 2005; Alonso-Nanclares et al., 2008; Blázquez-Llorca et al., 2010; Benavides-Piccione et al., 2013; Fernández-González et al., 2016; Peng et al., 2019; Montero-Crespo et al., 2020; Domínguez-Álvaro et al., 2021a; Cano-Astorga et al., 2021). In the present study, we have observed that interindividual variability may be related to certain cortical regions and to particular synaptic characteristics — with some regions more variable with regard to the synaptic density (e.g., dBA38), and others more variable with regard to the postsynaptic target distribution (e.g., BA21). Thus, similar or different synaptic characteristics can be found among individuals. The differences between the cortical circuit organization found in the present study between different individuals may suggest adaptations of each individual to particular functions.

## Material and Methods

### Tissue preparation

Human brain tissue was obtained from three autopsies (with short postmortem delays of less than 4 h) obtained from two men (53 and 66 years old) and one woman (53 years old) with no recorded neurological or psychiatric alterations (supplied by Unidad Asociada Neuromax, Laboratorio de Neuroanatomía Humana, Facultad de Medicina, Universidad de Castilla-La Mancha, Albacete). The procedure was approved by the Institutional Ethical Committee. This human brain tissue has been used in previous studies (Domínguez-Álvaro et al., 2018, 2019, 2021a; Montero-Crespo et al., 2020; Cano-Astorga et al., 2021).

Regarding the particular location of the sampled cortical areas, and in order to make a comprehensive correlation with the relevant literature, our samples from BA24 correspond to the “p24b” field defined in Palomero-Gallagher et al. (2008). BA38 has been distinguished at cytoarchitectonic (Ding et al., 2009; Blaizot et al., 2010; Insausti, 2013), connectivity (Kondo et al., 2003) and functional (Pascual et al., 2015) levels into many subdivisions. Thus, we have differentiated ventral BA38 (vBA38), which corresponds to the “TG” area defined in Ding et al. (2009); and dorsal BA38 (dBA38) which corresponds to the “TAr” area defined in Ding et al. (2009). BA21 corresponds to the middle temporal gyrus (T2).

After extraction, brain tissue was fixed in cold 4% paraformaldehyde (Sigma-Aldrich, St Louis, MO, USA) in 0.1 M sodium phosphate buffer (PB; Panreac, 131965, Spain), pH 7.4 for 24–48 h. After fixation, blocks of tissue of approximately 1cm x 1 cm x 1cm were washed in PB and sectioned coronally in a vibratome (150 μm thickness; Vibratome Sectioning System, VT1200S Vibratome, Leica Biosystems, Germany). Sections containing BA24, vBA38 and dBA38 from case AB7 were immersed in an antifreeze solution containing 30% glycerol (Panreac, 141339, Spain) and 30% ethylene glycol (Panreac, 141316, Spain) in 0.1M PB. Sections containing BA24, vBA38, dBA38 and BA21 from the rest of the cases were selected and processed for Nissl staining to determine cytoarchitecture (Fig. 5).

**Figure 5.**
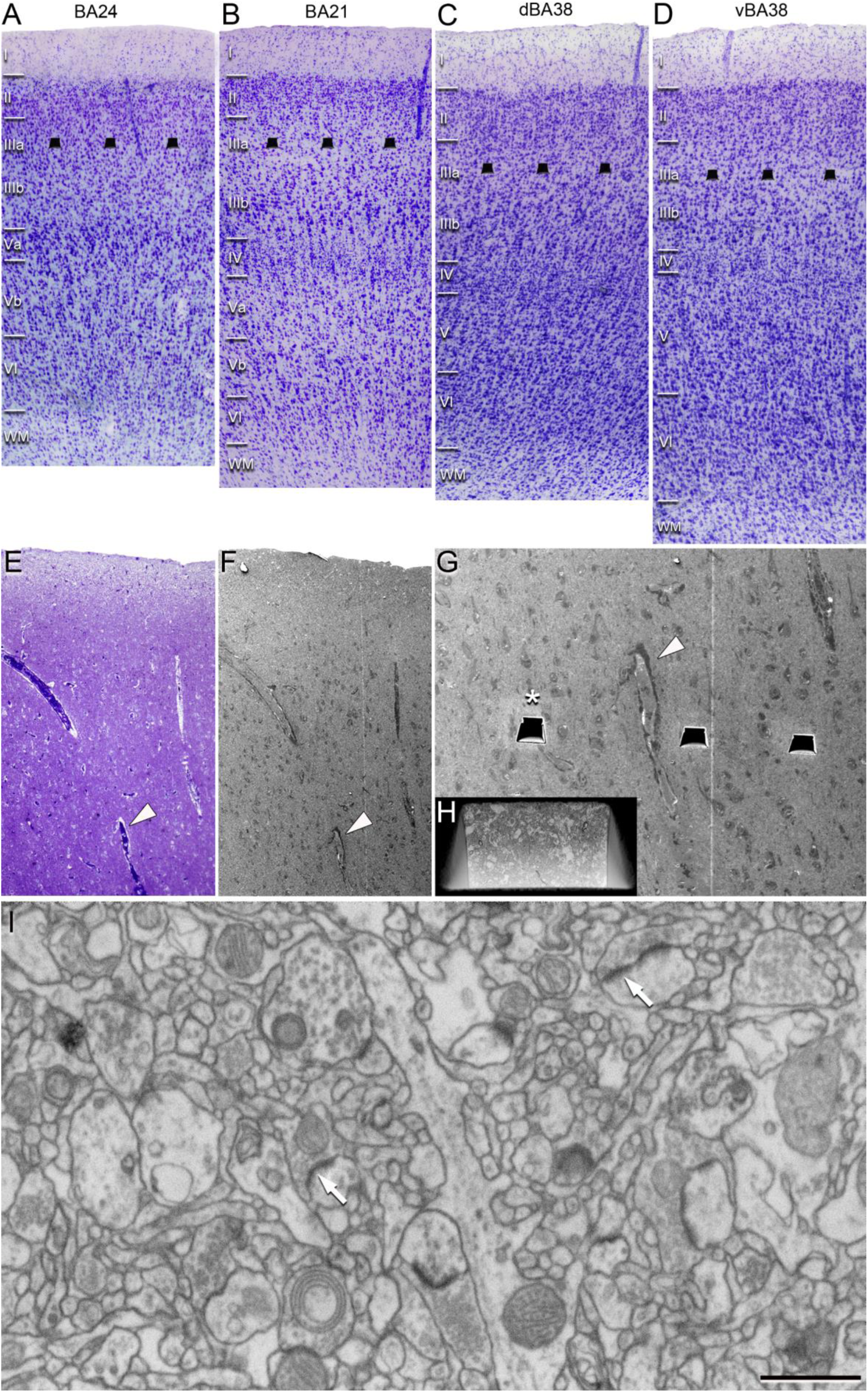
Cortical and FIB/SEM sampling regions. (A-D) Nissl-stained sections to illustrate the cytoarchitectonic differences between regions from an autopsy case. Cortical layer delimitation based on Palomero-Gallagher et al. (2008) for A, Alonso-Nanclares et al. (2008) in B, and Ding et al. (2009) for C and D. Illustration of the analyzed FIB/SEM sampling regions (superimposed as dark trapezoids in A-D). (E-G) Correlative light/electron microscopy analyses of layer III neuropil. (E) 1μm-thick semithin section stained with toluidine blue, which is adjacent to the block for FIB/SEM imaging (F). (F) SEM image illustrating the block surface. (G) Higher magnification of F, showing the trenches made in the neuropil to acquire the FIB/SEM stacks of images. White arrowheads in E, F and G point to the same blood vessel, allowing the exact location of the region of interest to be identified. (H) SEM image at higher magnification showing the front of a trench (white asterisk in G) made to acquire an FIB/SEM stack of images. (I) FIB/SEM image from a stack of images. Two synapses are indicated (arrows). Scale bar (in D) indicates 320 μm for A-D; 200 μm for E-F; 80 μm in G; 15 μm in H; 1.20 μm in I. BA: Brodmann’s area; d: dorsal; v: ventral.

### Volume fraction estimation of cortical elements

Semithin sections (1–1.5 μm thick) from all cases stained with toluidine blue (see below) were used to estimate the respective volume fraction (Vv) occupied by (i) neuropil, (ii) cell bodies (from neurons, glia and undetermined somata) and (iii) blood vessels. This estimation was performed applying the Cavalieri principle to 12 semithin sections per case (Gundersen et al., 1988) by point counting (Q-) using the integrated Stereo Investigator stereological package (Version 8.0, MicroBrightField Inc., VT, USA) attached to an Olympus light microscope (Olympus, Bellerup, Denmark) at 40× magnification. A grid, whose points covered an area of 2500 μm^2^, was randomly placed at six sites over the traced layer III on each semithin section to determine the Vv occupied by the different elements: neuropil, cell bodies and blood vessels (SI Fig. 1). Vv (in the case of the neuropil, for instance) was estimated with the following formula: Vv-neuropil = Q-neuropil *100 / (Q-neuropil + Q-neurons + Q-glia + Q-undetermined cells + Q-blood vessels).

### Electron Microscopy Processing

Sections containing BA24, vBA38, dBA38, and BA21 were selected, washed in 0.1M PB and postfixed for 24 h in a solution containing 2% paraformaldehyde, 2.5% glutaraldehyde (TAAB, G002, UK), and 0.003% CaCl_2_ (Sigma, C-2661-500G, Germany) in sodium cacodylate (Sigma, C0250-500G, Germany) buffer (0.1 M). The sections were treated with 1% OsO4 (Sigma, O5500, Germany), 0.1% potassium ferrocyanide (Probus, 23345, Spain), and 0.003% CaCl_2_ in sodium cacodylate buffer (0.1 M) for 1 h at room temperature. They were then stained with 1% uranyl acetate (EMS, 8473, USA), dehydrated, and flat-embedded in Araldite (TAAB, E021, UK) for 48 h at 60°C (DeFelipe and Fairén, 1993). The embedded sections were then glued onto a blank Araldite block. Semithin sections (1–2 μm thick) were obtained from the surface of the block and stained with 1% toluidine blue (Merck, 115930, Germany) in 1% sodium borate (Panreac, 141644, Spain). The last semithin section (which corresponds to the section immediately adjacent to the block surface) was examined under light microscope and photographed to accurately locate the neuropil regions to be examined (Fig. 5).

### Three-Dimensional Electron Microscopy

The 3D study of the samples was carried out using a dual beam microscope (Crossbeam® 40 electron microscope, Carl Zeiss NTS GmbH, Oberkochen, Germany). This instrument combines a high-resolution field-emission SEM column with a focused gallium ion beam (FIB), which permits removal of thin layers of material from the sample surface on a nanometer scale. As soon as one layer of material (20 nm thick) is removed by the FIB, the exposed surface of the sample is imaged by the SEM using the backscattered electron detector. The sequential automated use of FIB milling and SEM imaging allowed us to obtain long series of photographs of a 3D sample of selected regions (Merchán-Pérez et al., 2009). Image resolution in the xy plane was 5 nm/pixel. Resolution in the z-axis (section thickness) was 20 nm, and image size was 2048×1536 pixels. These parameters were chosen in order to obtain a large enough field of view where synaptic junctions could be clearly identified, within a reasonable time frame (approximately 12 h per stack of images). The number of sections per stack ranged from 240–330 in BA24, 254–314 in vBA38, 261–305 in dBA38, and 260–317 in BA21, which corresponds to a volume per stack ranging from 623–856 μm^3^ (mean: 709 μm^3^) in BA24, 654–815 μm^3^ (mean: 726 μm^3^) in vBA38, 678–792 μm^3^ (mean: 716 μm^3^) in dBA38, and 675–822 μm^3^ (mean: 744 μm^3^) in BA21. We acquired 9 stacks of images (layer III; approximately 600–1,100 μm from the pial surface) in BA24 (three stacks per case, in three cases; total volume studied: 4289 μm^3^); 9 stacks of images (approximately 700–800 μm from the pial surface) in vBA38 (three stacks per case, in three cases; total volume studied: 4,399 μm^3^); 9 stacks of images (approximately 600–900 μm from the pial surface) in dBA38 (three stacks from three cases; total volume studied: 4,336 μm^3^); and 7 stacks of images (approximately 600 μm from the pial surface) in BA21 (two stacks of images per case in cases AB2 and AB3 taken from Cano-Astorga et al. (2021), and three stack of images in case AB7; total volume studied: 3,896 μm^3^). Examples of the serial images obtained in each cortical region are illustrated in SI Fig. 8.

All measurements were corrected for tissue shrinkage, which occurs during the processing of sections (Merchán-Pérez et al., 2009). To estimate the shrinkage in our samples, we photographed and measured the area of the vibratome sections with ImageJ (ImageJ 1.51; NIH, USA), both before and after processing for electron microscopy. The section area values after processing were divided by the values before processing to obtain the volume, area, and linear shrinkage factors (Oorschot et al., 1991) — yielding correction factors of 0.90, 0.93, and 0.97, respectively. Nevertheless, in order to compare with previous studies —in which either no correction factors had been included or such factors were estimated using other methods— in the present study, we provided both sets of data. Additionally, a correction in the volume of the stack of images —to account for the presence of fixation artifact (i.e., swollen neuronal or glial processes)— was applied after quantification with Cavalieri principle (Gundersen et al., 1988; see Montero-Crespo et al., 2020). Every FIB/SEM stack was examined, and the volume artifact was found to range from 0.0 to 16.1% of the volume stacks.

### Three-Dimensional Analysis of Synapses

Stacks of images obtained by the FIB/SEM were analyzed using EspINA software (EspINA Interactive Neuron Analyzer, 2.9.12; https://cajalbbp.es/espina/; Morales et al., 2011; Fig. 6). As previously discussed (Merchán-Pérez et al., 2009), there is a consensus for classifying cortical synapses into asymmetric synapses (AS; or type I) and symmetric synapses (SS; or type II). The main characteristic distinguishing these synapses is their prominent or thin postsynaptic density, respectively (SI Fig. 9). Also, these two types of synapses correlate with different functions: AS are mostly glutamatergic and excitatory, while SS are mostly GABAergic and inhibitory (DeFelipe and Fariñas, 1992; DeFelipe et al., 1999). Nevertheless, in single sections, the synaptic cleft and the pre- and postsynaptic densities are often blurred if the plane of the section does not pass at right angles to the synaptic junction. Since the software EspINA allows navigation through the stack of images, it was possible to unambiguously identify every synapse as AS or SS based on the thickness of the postsynaptic density (PSD) (Merchán-Pérez et al., 2009).

**Figure 6.**
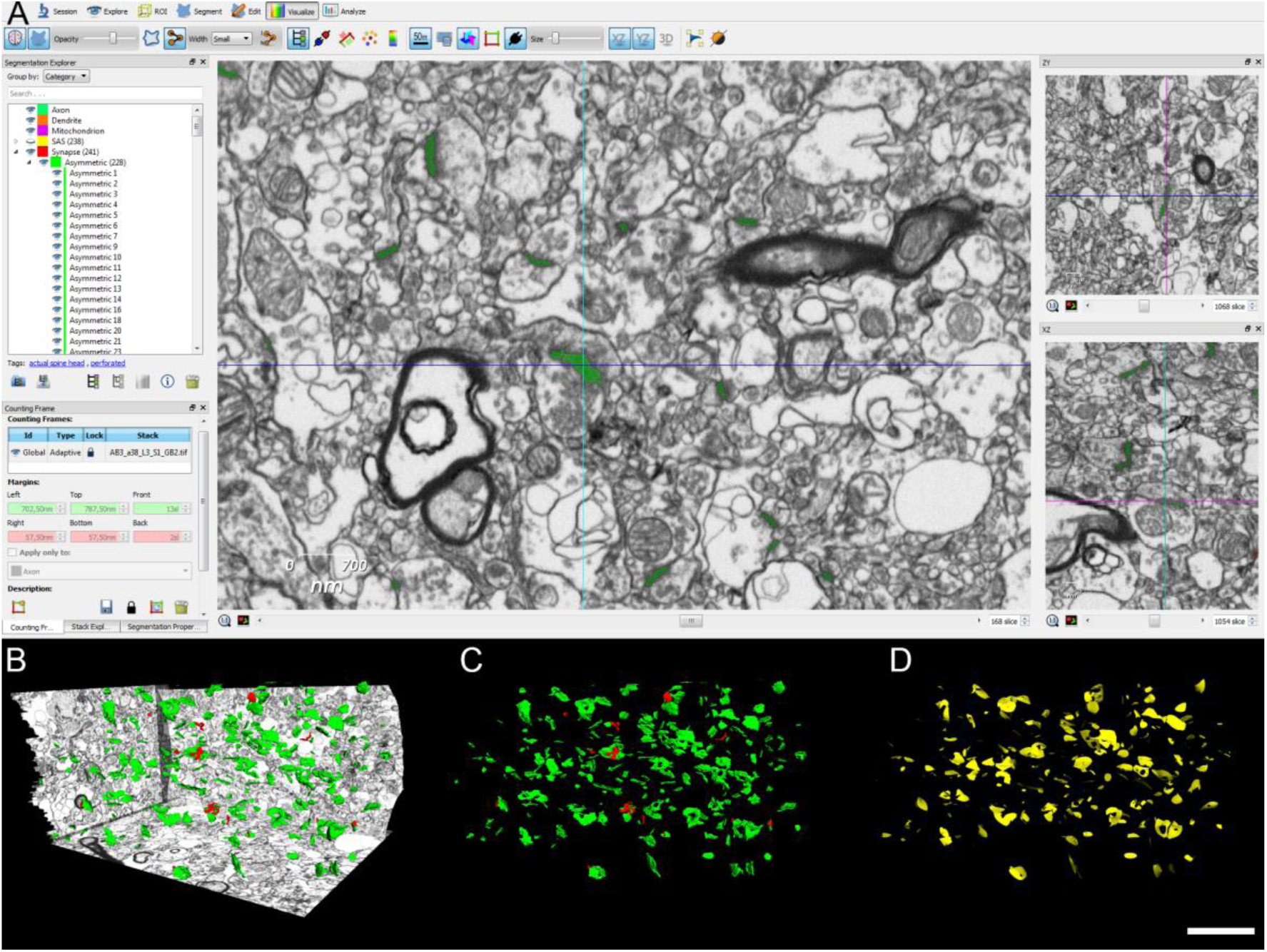
Identification and annotation of synapses. (A-D) Screenshots of the EspINA software user interface. (A) In the main window, the sections are viewed through the xy plane (as obtained by FIB/SEM microscopy). The other two orthogonal planes, yz and xz, are also shown in adjacent windows (on the right). (B) The 3D window shows the three orthogonal planes and the 3D reconstruction of asymmetric (green) and symmetric (red) synapses. (C) 3D Reconstructed synapses displayed by colors. (D) Computed Synaptic Apposition Surface for each reconstructed synapse (yellow). Scale bar (in D) indicates 2600 nm, for B–D.

Synapses with prominent PSDs are classified as AS, while those with thin PSDs are classified as SS (Gray, 1959; Peters et al., 1991; Fig. 5). In addition, geometrical features —such as size and shape— and spatial distribution features (centroids) of each reconstructed synapse were also calculated by EspINA. This software also extracts the Synaptic Apposition Area (SAS) and provides its measurements (Fig. 6). Given that the pre- and postsynaptic densities are located face to face, their surface areas are comparable (for details, see Morales et al., 2013). Since the SAS comprises both the active zone and the PSD, it is a functionally relevant measure of the size of a synapse (Morales et al., 2013). EspINA was also used to visualize each of the reconstructed synapses in 3D and to detect the possible presence of perforations or deep indentations in their perimeters. Regarding the shape of the PSD, the synaptic junctions could be classified into four main categories, according to the categories proposed by Santuy et al. (2018a): macular (disk-shaped PSD), perforated (with one or more holes in the PSD), horseshoe-shaped (with an indentation), and fragmented (disk-shaped PSDs with no connection between them) (Fig. 2C). To identify the postsynaptic targets of the synapses, we navigated through the image stack using EspINA to determine whether the postsynaptic element was a dendritic spine (spine, for simplicity) or a dendritic shaft. As previously described in Domínguez-Álvaro et al. (2021a, 2021b), unambiguous identification of spines requires the spine to be visually traced to the parent dendrite (see Cano-Astorga et al., 2021), in which case we refer to them as complete spines. When synapses were established on a spine head-shaped postsynaptic element whose neck could not be followed to the parent dendrite, we identified these elements as incomplete spines. These incomplete spines were identified on the basis of their size and shape, the lack of mitochondria, and the presence of a spine apparatus — or because they were filled with ‘fluffy material’ (a term coined by Peters et al. (1991) to describe the fine and indistinct filaments present in the spines; see also del Río and DeFelipe, 1995). For simplicity, we will refer to both the complete and incomplete spines as spines, unless otherwise specified. We also recorded the presence of single or multiple synapses on a single spine. Furthermore, we determined whether the target dendrite had spines or not.

### Quantification of the Synaptic Density

EspINA provided the 3D reconstruction of every synapse and allowed the application of an unbiased 3D counting frame (CF) — a regular prism enclosed by three acceptance planes and three exclusion planes marking its boundaries. All objects within the CF are counted, as are those intersecting any of the acceptance planes, while objects that are outside the CF, or intersecting any of the exclusion planes, are not counted. Thus, the number of synapses per unit volume was calculated directly by dividing the total number of synapses counted by the volume of the CF (Merchán-Pérez et al., 2009). This method was used in all 37 stacks of images (three stacks of images per case, in three cases in BA24, vBA38, dBA38; two stacks of images per case in cases AB2 and AB3, taken from Cano-Astorga et al. (2021), and three stacks of images in case AB7 in BA21.

### Spatial Distribution Analysis of Synapses

To analyze the spatial distribution of synapses, spatial point pattern analysis was performed as described elsewhere (Antón-Sánchez et al., 2014; Merchán-Pérez et al., 2014). Briefly, we compared the actual position of centroids of synapses with the CSR model — a random spatial distribution model that defines a situation where a point is equally likely to occur at any location within a given volume. For each of the 37 FIB/SEM stacks of images, we calculated three functions commonly used for spatial point pattern analysis: G, F, and K functions (SI Fig. 2). As described in Merchán-Pérez et al. (2014) (see also Antón-Sánchez et al., 2014), the G function, also called the nearest-neighbor distance cumulative distribution function or the event-to-event distribution, is —for a distance d— the probability that a typical point separates from its nearest neighbor by a distance of d at the most. The F function, also known as the empty space function or the point-to-event distribution, is —for a distance d— the probability that the distance of each point (in a regularly spaced grid of L points superimposed over the sample) to its nearest synapse centroid is d at the most. The K function, also called the reduced second moment function or Ripley’s function, is —for a distance d— the expected number of points within a distance d of a typical point of the process divided by the intensity λ. An estimation of the K function is given by the mean number of points within a sphere of increasing radius d centered on each sample point, divided by an estimation of the expected number of points per unit volume. This study was carried out using the Spatstat package and R Project program (Baddeley et al., 2015).

### Statistical Analysis

One-way analyses of variance (ANOVA) with Holm-Sidak’s post-hoc correction was performed to compare the synaptic density between regions. To perform statistical comparisons of AS:SS proportions regarding their synaptic shape and their postsynaptic targets, chi-square (χ^2^) test was used for contingency tables. The same method was used to obtain comparisons between regions with regard to the volume occupied by the cortical structures, the synaptic type, the synaptic shapes, and their postsynaptic targets. Kruskal– Wallis (KW) nonparametric test, with post-hoc correction (via Dunn’s multiple comparisons) was performed to analyze the area of the SAS. Frequency distribution analysis of the SAS was performed using Kolmogorov–Smirnov (KS) nonparametric test.

Statistical studies were performed with the GraphPad Prism statistical package (Prism 9.00 for Windows, GraphPad Software Inc., USA), Spatstat package and R Project program (Baddeley et al., 2015), as well as the on-line tool VassarStats (http://vassarstats.net/).

### Ethics approval

Brain tissue samples were obtained following the guidelines and approval of the Institutional Ethical Committee at the School of Medicine, University of Castilla-La Mancha (Albacete, Spain).

## Abbreviation list

3D: three-dimensional
AMPA: α-amino-3-hydroxy-5-methyl-4-isoxazolepropionic acid
AS: asymmetric synapses
CA1: Cornu Ammonis 1
CF: counting frame
CSR: Complete Spatial Randomness
FIB/SEM: focused ion beam/scanning electron microscopy
KS: Kolmogorov-Smirnov
MW: Mann-Whitney
NMDA: N-methyl-D-aspartate
PB: phosphate buffer
PSD: postsynaptic density
SAS: synaptic apposition surface
SE: standard error of the mean
SD: standard deviation
SS: symmetric synapses
Vv: volume fraction.

## Availability of data and materials

Most data are available in the main text and the SI. The datasets used and analyzed during the current study are published in the EBRAINS Knowledge Graph (DOI: 10.25493/B3V0-4D8): Alonso-Nanclares L., Cano-Astorga N., Plaza-Alonso S., DeFelipe J. (2022). “3D ultrastructural study of synapses using FIB/SEM in the Human Cortex (Brodmann’s areas 24 and 38)”. Human Brain Project Neuroinformatics Platform.

## Acknowledgements

This work was supported by the following Grants: PGC2018-094307-B-I00 (to J.D.) funded by MCIN/AEI/10.13039/501100011033, CSIC Interdisciplinary Thematic Platform - Cajal Blue Brain (PTI-BLUEBRAIN; Spain) and the European Union’s Horizon 2020 Framework Programme for Research and Innovation under Specific Grant Agreement No. 945539 (Human Brain Project SGA3). Research Fellowships funded by MCIN/AEI/10.13039/501100011033 for N.C.-A. (PRE2019-089228) and S.P.-A. (FPU19/00007).

We would like to thank Carmen Álvarez and Lorena Valdés for their technical assistance, and Nick Guthrie for his excellent editorial assistance.

