## Supplementary material for "3D Synaptic Organization of Layer III of the Human Anterior Cingulate and Temporopolar Cortex": SI Table 1

**Supplementary Information**

| Cortical area | Case | $V_v$ Neuropil | $V_v$ Cell Bodies | $V_v$ Blood Vessels |
|---|---|---|---|---|
| **BA24** | AB2 | 82.76% (3,393) | 11.32% (464) | 5.93% (243) |
| | AB3 | 86.72% (2,515) | 11.41% (331) | 1.86% (54) |
| | AB7 | 83.63% (2,676) | 12.91% (413) | 3.47% (111) |
| | *All* | *84.37% (8584)* | *11.88% (1,208)* | *3.75% (408)* |
| **vBA38** | AB2 | 85.19% (2,641) | 10.32% (320) | 4.48% (139) |
| | AB3 | 92.71% (2,772) | 3.61% (108) | 3.68% (110) |
| | AB7 | 91.23% (2,828) | 6.45% (200) | 2.32% (72) |
| | *All* | *89.71% (8,241)* | *6.80% (628)* | *3.50% (321)* |
| **dBA38** | AB2 | 85.89% (3,865) | 10.69% (481) | 3.42% (154) |
| | AB3 | 90.14% (2,524) | 7.43% (208) | 2.43% (68) |
| | AB7 | 90.65% (4,714) | 6.47% (321) | 3.17% (165) |
| | *All* | *88.90% (11,103)* | *8.09% (1,010)* | *3.01% (387)* |
| **BA21** | AB2 | 88.31% (2,296) | 9.85% (256) | 1.85% (48) |
| | AB3 | 90.26% (2,076) | 6.87% (158) | 2.87% (66) |
| | AB7 | 86.71% (3,295) | 9.58% (364 ) | 3.71% (141) |
| | *All* | *88.43% (7,667)* | *8.76% (778)* | *2.81% (255)* |

**SI Table 1.** Volume fraction ($V_v$) occupied by different cortical elements in BA24, vBA38, dBA38 and BA21 per case. The absolute numbers below the percentages of volume fractions indicate the number of times that a grid point was counted in each category (neuropil, cell bodies and blood vessels). BA: Brodmann’s area; d: dorsal; v: ventral.

| Cortical area | | **BA24** | | | **vBA38** | | | **dBA38** | | | **BA21** | |
|---|---|---|---|---|---|---|---|---|---|---|---|---|
| Case | AB2 | AB3 | AB7 | AB2 | AB3 | AB7 | AB2 | AB3 | AB7 | AB2 | AB3 | AB7 |
| No. AS | 399 | 420 | 516 | 632 | 477 | 435 | 688 | 590 | 1001 | 372 | 307 | 450 |
| No. SS | 33 | 37 | 25 | 27 | 30 | 28 | 55 | 34 | 56 | 23 | 30 | 30 |
| No. synapses (AS+SS) | 432 | 457 | 541 | 659 | 507 | 463 | 743 | 624 | 1057 | 395 | 337 | 480 |
| % AS | 92.36±0.53 | 91.89±1.44 | 95.38±1.68 | 95.90±0.65 | 94.08±1.29 | 93.95±1.77 | 92.60±2.21 | 94.55±2.90 | 94.70±1.16 | 94.13±4.09 | 91.04±1.55 | 93.67±2.19 |
| % SS | 7.64±0.53 | 8.11±1.44 | 4.62±1.68 | 4.10±0.65 | 5.92±1.29 | 6.05±1.77 | 7.40±2.21 | 5.45±2.90 | 5.30±1.16 | 5.87±4.09 | 8.96±1.55 | 6.33±2.19 |
| CF volume ($\mu m^3$) | 955 (860) | 1,050 (946) | 1,139 (1,025) | 1,065 (959) | 1,058 (953) | 987 (889) | 1,014 (913) | 958 (863) | 1,194 (1,075) | 794 (715) | 764 (688) | 925 (833) |
| No. AS/$\mu m^3$ (mean ± SD) | 0.52±0.03 (0.57±0.04) | 0.40±0.02 (0.44±0.03) | 0.45±0.02 (0.50±0.02) | 0.59±0.05 (0.66±0.05) | 0.45±0.02 (0.50±0.03) | 0.44±0.06 (0.49±0.06) | 0.68±0.14 (0.75±0.16) | 0.62±0.06 (0.68±0.07) | 0.84±0.04 (0.93±0.04) | 0.47±0.06 (0.52±0.07) | 0.40±0.03 (0.45±0.03) | 0.48±0.08 (0.53±0.08) |
| No. SS/$\mu m^3$ (mean ± SD) | 0.03±0.00 (0.04±0.01) | 0.04±0.01 (0.04±0.01) | 0.02±0.01 (0.03±0.01) | 0.03±0.01 (0.03±0.01) | 0.03±0.01 (0.03±0.01) | 0.03±0.01 (0.03±0.01) | 0.05±0.02 (0.06±0.02) | 0.04±0.02 (0.04±0.02) | 0.05±0.01 (0.05±0.01) | 0.03±0.02 (0.03±0.02) | 0.04±0.00 (0.04±0.01) | 0.03±0.01 (0.04±0.01) |
| No. all synapses/$\mu m^3$ (mean ± SD) | 0.55±0.03 (0.61±0.04) | 0.43±0.02 (0.48±0.02) | 0.48±0.03 (0.53±0.03) | 0.62±0.05 (0.69±0.06) | 0.48±0.02 (0.53±0.02) | 0.67±0.06 (0.52±0.07) | 0.73±0.16 (0.81±0.17) | 0.65±0.07 (0.72±0.08) | 0.89±0.03 (0.98±0.03) | 0.50±0.05 (0.55±0.05) | 0.44±0.02 (0.49±0.03) | 0.51±0.08 (0.57±0.09) |
| Intersynaptic distance (nm; mean ± SD) | 814±91 (757±84) | 856±8 (796±8) | 864±19 (802±18) | 789±38 (733±35) | 834±16 (775±15) | 842±47 (782±44) | 753±34 (700±32) | 778±28 (723±26) | 685±15 (637±14) | 760±86 (737±83) | 857±94 (831±91) | 839±99 (814±96) |
| Area of SAS AS ($nm^2$; mean ± SE) | 109,290± 4,148 (101,640± 3,857) | 151,803± 4,854 (141,176± 4,514) | 101,096± 3,324 (94,019± 3,091) | 111,306± 3,401 (103,514± 3,163) | 125,118± 4,805 (116,360± 4,468) | 96,187± 3,669 (102,094± 3,008) | 95,178± 3,189 (88,515± 2,966) | 79,510± 3,095 (73,945± 2,878) | 63,663± 1,812 (59,176± 1,685) | 116,131± 4,473 (108,002± 4,160) | 132,104± 5,626 (122,857± 5,232) | 105,541± 3,988 (98,153± 3,709) |
| Area of SAS SS ($nm^2$; mean ± SE) | 51,396± 6,294 (47,798± 5,854) | 89,353± 11,848 (83,098± 11,019) | 56,557± 6,355 (52,598± 5,910) | 58,660± 7,641 (54,554± 7,106) | 61,383± 9,305 (57,086± 8,654) | 48,704± 5,017 (45,294± 4,665) | 58,982± 4,735 (54,853± 4,403) | 53,343± 4387 (49,609± 4,080) | 45,519± 4,327 (42,333± 4,024) | 57,193± 9,384 (53,190± 8,728) | 84,027± 1,0474 (78,145± 9,741) | 45,833± 5,908 (42,625± 5,494) |

**SI Table 2.** Accumulated data obtained from the ultrastructural analysis of neuropil from layer III of BA24, vBA38, dBA38 and BA21 per case. Data in parentheses have not been corrected for shrinkage. AS: asymmetric synapses; BA: Brodmann's area; CF: counting frame; d: dorsal; SAS: synaptic apposition surface; SD: standard deviation; SE: standard error of the mean; SS: symmetric synapses; v: ventral.

| Cortical area | Case | Type of synapse | Macular synapses | Perforated synapses | Horseshoe-shaped synapses | Fragmented synapses | Total synapses |
|---|---|---|---|---|---|---|---|
| **BA24** | AB2 | AS | 85.2% (340) | 9.0% (36) | 5.0% (20) | 0.8% (3) | 100% (399) |
| | | SS | 84.8% (28) | 3.0% (1) | 12.1% (4) | 0.0% (0) | 100% (33) |
| | AB3 | AS | 75.5% (317) | 16.2% (68) | 7.1% (30) | 1.2% (5) | 100% (420) |
| | | SS | 83.8% (31) | 13.5% (5) | 2.7% (1) | 0.0% (0) | 100% (37) |
| | AB7 | AS | 82.6% (426) | 10.1% (52) | 5.4% (28) | 1.9% (10) | 100% (516) |
| | | SS | 88.0% (22) | 8.0% (2) | 4.0% (1) | 0.0% (0) | 100% (25) |
| | *All* | *AS* | *81.2% (1083)* | *11.7% (156)* | *5.8% (78)* | *1.3% (18)* | *100% (1,335)* |
| | | *SS* | *85.3% (81)* | *8.4% (8)* | *6.3% (6)* | *0% (0)* | *100% (95)* |
| **vBA38** | AB2 | AS | 81.2% (513) | 12.3% (78) | 5.4% (34) | 1.1% (7) | 100% (632) |
| | | SS | 85.2% (23) | 3.7% (1) | 3.7% (1) | 7.4% (2) | 100% (27) |
| | AB3 | AS | 80.3% (383) | 14.5% (69) | 2.9% (14) | 2.3% (11) | 100% (477) |
| | | SS | 100.0% (30) | 0.0% (0) | 0.0% (0) | 0.0% (0) | 100% (30) |
| | AB7 | AS | 85.7% (373) | 6.4% (28) | 4.6% (20) | 3.0% (13) | 100% (435) |
| | | SS | 85.7% (24) | 7.1% (2) | 3.6% (1) | 3.6% (1) | 100% (28) |
| | *All* | *AS* | *82.2% (1269)* | *11.3% (175)* | *4.4% (68)* | *2.1% (31)* | *100% (1,586)* |
| | | *SS* | *90.6% (77)* | *3.5% (3)* | *2.4% (2)* | *3.5% (3)* | *100% (85)* |
| **dBA38** | AB2 | AS | 86.8% (597) | 10.2% (70) | 2.3% (16) | 0.7% (5) | 100% (688) |
| | | SS | 90.9% (50) | 3.6% (2) | 3.6% (2) | 1.8% (1) | 100% (55) |
| | AB3 | AS | 83.2% (491) | 9.8% (58) | 4.4% (26) | 2.2% (13) | 100% (590) |
| | | SS | 94.1% (32) | 2.9% (1) | 2.9% (1) | 0.0% (0) | 100% (34) |

| | | | | | | | |
|---|---|---|---|---|---|---|---|
| | AB7 | AS | 82.8% (829) | 10.1% (101) | 5.3% (53) | 1.8% (18) | 100% (1,001) |
| | | SS | 91.1% (51) | 1.8% (1) | 7.1% (4) | 0.0% (0) | 100% (56) |
| | *All* | *AS* | *84.1% (1917)* | *10.1% (229)* | *4.2% (95)* | *1.6% (36)* | *100% (2,279)* |
| | | *SS* | *91.7% (133)* | *2.8% (4)* | *4.8% (7)* | *0.7% (1)* | *100% (145)* |
| **BA21** | AB2 | AS | 77.2% (287) | 18.3% (68) | 3.2% (12) | 1.3% (5) | 100% (372) |
| | | SS | 95.7% (22) | 4.3% (1) | 0.0% (0) | 0.0% (0) | 100% (23) |
| | AB3 | AS | 75.9% (233) | 19.5% (60) | 3.9% (12) | 0.7% (2) | 100% (307) |
| | | SS | 80.0% (24) | 3.3% (1) | 16.7% (5) | 0.0% (0) | 100% (30) |
| | AB7 | AS | 77.6% (349) | 14.9% (67) | 5.6% (25) | 2.0% (9) | 100% (450) |
| | | SS | 96.7% (29) | 0.0% (0) | 0.0% (0) | 3.3% (1) | 100% (30) |
| | *All* | *AS* | *77.0% (869)* | *17.3% (195)* | *4.3% (49)* | *1.4% (16)* | *100% (1,129)* |
| | | *SS* | *90.4% (75)* | *2.4% (2)* | *6.0% (5)* | *1.2% (1)* | *100% (83)* |

**SI Table 3.** Proportions of the different synaptic shapes from layer III of BA24, vBA38, dBA38 and BA21 per case. Data are given as percentages (absolute numbers of synapses studied are given in parentheses). AS: asymmetric synapses; BA: Brodmann's area; d: dorsal; SS: symmetric synapses; v: ventral.

| Cortical area | Type of synapses | Morphological category | Area of SAS (mean ± SE) |
|---|---|---|---|
| BA24 | AS | Macular | 92,550±1,831 (86,071±1,703) |
| | | Perforated | 244,324±8,586 (227,222±7,985) |
| | | Horseshoe | 223,314±8,573 (207,682±7,973) |
| | | Fragmented | 213,303±19,351 (198,372±17,996) |
| | SS | Macular | 57,133±2,489 (53,134±4,341) |
| | | Perforated | 159,819±31,934 (148,632±29,669) |
| | | Horseshoe | 84,954±13,223 (79,007±12,297) |
| | | Fragmented | NA |
| vBA38 | AS | Macular | 81,046±1,468 (75,373±1,366) |
| | | Perforated | 255,828±8,012 (237,920±7,451) |
| | | Horseshoe | 235,593±8,399 (219,101±7,811) |
| | | Fragmented | 258,032±16,807 (239,970±15,631) |
| | SS | Macular | 56,222±4,204 (49,895±3,898) |
| | | Perforated | 75,745±21,752 (70,443±20,229) |
| | | Horseshoe | 79,056±22,726 (73,522±21,135) |
| | | Fragmented | 63,564±10,331 (59,115±9,608) |
| dBA38 | AS | Macular | 56,222±977(52,286±909) |
| | | Perforated | 191,842±6,389 (178,413±5,942) |
| | | Horseshoe | 180,592±8,095 (167,951±7,528) |
| | | Fragmented | 187,816±11,500 (183,969±10,695) |
| | SS | Macular | 50,487±2,838 (47,067±2,500) |
| | | Perforated | 66,017±13,889 (61,396±12,917) |
| | | Horseshoe | 84,193±17,302 (78,229±16,091) |
| | | Fragmented | 22,212±NA (20,657±NA) |
| BA21 | AS | Macular | 79,659±1,460 (74,082±1,358) |
| | | Perforated | 218,035±6,714 (202,773±6,244) |
| | | Horseshoe | 214,517±12,230 (199,501±11,374) |
| | | Fragmented | 252,169±12,163 (234,517±11,312) |
| | SS | Macular | 66,412±4,746 (61,763±4,413) |
| | | Perforated | 106,608±30,659 (9,9145±28,513) |

| | | |
|---|---|---|
| | Horseshoe | 87,519±25,992 (81,393±24,173) |
| | Fragmented | 69,672±16,385 (64,795±15,238) |

**SI Table 4.** Area of the SAS ($nm^2$) of macular, perforated, horseshoe and fragmented AS and SS in layer III of BA24, vBA38, dBA38 and BA21. SAS areas of the macular AS were significantly smaller than perforated, horseshoe and fragmented in all regions (KW, $p<0.0001$). Data in parentheses are not corrected for shrinkage. AS: asymmetric synapses; BA: Brodmann's area; d: dorsal; NA: not applicable; SAS: synaptic apposition surface; SE: standard error of the mean; SS: symmetric synapses; v: ventral.

| Cortical area | Case | Type of synapse | Synapses on complete spines | Synapses on incomplete spines | Synapses on spine necks | Synapses on dendritic shafts with spines | Synapses on dendritic shafts without spines | Total synapses |
|---|---|---|---|---|---|---|---|---|
| **BA24** | AB2 | AS | 28.66% (88) | 43.65% (134) | 0.98% (3) | 15.31% (47) | 11.40% (35) | 100% (307) |
| | | SS | 7.69% (2) | 7.69% (2) | 3.85% (1) | 42.31% (11) | 38.46% (10) | 100% (26) |
| | AB3 | AS | 50.77% (199) | 24.74% (97) | 1.02% (4) | 13.78% (54) | 9.69% (38) | 100% (392) |
| | | SS | 25.71% (9) | 8.57% (3) | 5.71% (2) | 37.14% (13) | 22.86% (8) | 100% (35) |
| | AB7 | AS | 51.46% (247) | 20.63% (99) | 1.04% (5) | 13.75% (66) | 13.13% (63) | 100% (480) |
| | | SS | 4.55% (1) | 0.00% (0) | 18.18% (4) | 18.18% (4) | 59.09% (13) | 100% (22) |
| | *All* | *AS* | *45.29% (534)* | *27.99 (330)* | *1.02% (12)* | *14.16% (167)* | *11.5 4% (136)* | *100% (1,179)* |
| | | *SS* | *14.46% (12)* | *6.02% (5)* | *8.43% (7)* | *33.73% (28)* | *37.35% (31)* | *100% (83)* |
| **vBA38** | AB2 | AS | 36.00% (216) | 41.83% (251) | 0.17% (1) | 9.83% (59) | 12.17% (73) | 100% (600) |
| | | SS | 7.69% (2) | 7.69% (2) | 3.85% (1) | 53.85% (14) | 26.92% (7) | 100% (26) |
| | AB3 | AS | 34.33% (160) | 45.92% (214) | 0.64% (3) | 10.09% (47) | 9.01% (42) | 100% (466) |
| | | SS | 4.14% (2) | 28.57% (8) | 7.14% (2) | 35.71% (10) | 21.43% (6) | 100% (28) |

| | | | | | | | | |
|---|---|---|---|---|---|---|---|---|
| | AB7 | AS | 50.70% (216) | 23.00% (98) | 1.17% (5) | 14.32% (61) | 10.80% (46) | 100% (426) |
| | | SS | 14.29% (4) | 3.57% (1) | 3.57% (1) | 60.71% (17) | 17.86% (5) | 100% (28) |
| | *All* | *AS* | *39.68% (592)* | *37.73% (563)* | *0.60% (9)* | *11.19% (167)* | *10.79% (161)* | *100% (1,492)* |
| | | *SS* | *9.76% (8)* | *13.41% (11)* | *4.88% (4)* | *50.00% (41)* | *21.95% (18)* | *100% (82)* |
| **dBA38** | AB2 | AS | 23.93 % (157) | 51.22% (336) | 0.30% (2) | 10.21% (67) | 14.33% (94) | 100% (656) |
| | | SS | 3.64% (2) | 7.27% (4) | 7.27% (4) | 29.09% (16) | 52.73% (29) | 100% (55) |
| | AB3 | AS | 30.33% (175) | 52.86% (305) | 0.69% (4) | 8.15% (47) | 7.97% (46) | 100% (577) |
| | | SS | 9.38% (3) | 18.75% (6) | 3.13% (1) | 50.00% (16) | 18.75% (6) | 100% (32) |
| | AB7 | AS | 48.87% (477) | 23.05% (225) | 1.23% (12) | 16.91% (165) | 9.94% (97) | 100% (976) |
| | | SS | 16.67% (9) | 3.70% (2) | 5.56% (3) | 55.56% (30) | 18.52% (10) | 100% (54) |
| | *All* | *AS* | *36.62% (809)* | *39.20% (866)* | *0.81% (18)* | *12.63% (279)* | *10.73% (237)* | *100% (2,209)* |
| | | *SS* | *9.93% (14)* | *8.51% (12)* | *5.67% (8)* | *43.97% (62)* | *31.91% (45)* | *100% (141)* |
| **BA21** | AB2 | AS | 47.3% (159) | 23.8% (80) | 0.9% (3) | 11.9% (40) | 16.1% (54) | 100% (336) |
| | | SS | 4.3% (1) | 4.3% (1) | 4.3% (1) | 52.2% (12) | 34.8% (8) | 100% (23) |

| | | | | | | | |
|---|---|---|---|---|---|---|---|
| AB3 | AS | 51.2% (144) | 16.0% (45) | 2.5% (7) | 23.1% (65) | 7.1% (20) | 100% (281) |
| | SS | 19.2% (5) | 3.8% (1) | 3.8% (1) | 42.3% (11) | 30.8% (8) | 100% (26) |
| AB7 | AS | 63.2% (283) | 18.3% (82) | 0.7% (3) | 6.7% (30) | 11.2% (50) | 100% (448) |
| | SS | 3.3% (1) | 3.3% (1) | 10% (3) | 43.3% (13) | 40.0% (12) | 100% (30) |
| *All* | *AS* | *55.02% (586)* | *19.44% (207)* | *1.22% (13)* | *12.68% (135)* | *11.64% (124)* | *100% (1,065)* |
| | *SS* | *7.69% (6)* | *3.85% (3)* | *5.13% (4)* | *47.44% (37)* | *335.90% (28)* | *100% (78)* |

**SI Table 5.** Distribution of AS and SS on spines and dendritic shafts in layer III of BA24, vBA38, dBA38 per case. Synapses on spines have been sub-divided into those that are established on spine heads and those established on spine necks. Data are given as percentages with the absolute number of synapses studied in parentheses. AS: asymmetric synapses; BA: Brodmann's area; d: dorsal; SS: symmetric synapses; v: ventral.

| Cortical area | Type of synapses | Postsynaptic target | Area of SAS (mean ± SE) |
|---|---|---|---|
| BA24 | AS | Complete spine heads | 134,358±4,109 (126,632±3,960) |
| | | Incomplete spine heads | 123,943±5,131 (113,180±5,298) |
| | | Spine necks | 70,559±10,598 (65,619±9,856) |
| | | Shafts with spines | 113,180±5,298 (105,257±4,927) |
| | | Shafts without spines | 101,700±6,050 (94,581±5,626) |
| | SS | Complete spine heads | 99,812±27,941 (92,826±25,985) |
| | | Incomplete spine heads | 42,617±21,129 (39,634±19,650) |
| | | Spine necks | 56,914±9,504 (52,930±8,839) |
| | | Shafts with spines | 76,834±11,512 (71,455±10,706) |
| | | Shafts without spines | 58,578±6,408 (54,478±5,960) |
| vBA38 | AS | Complete spine heads | 131,073±4,241 (121,898±3,944) |
| | | Incomplete spine heads | 106,156±3,806 (98,725±3,540) |
| | | Spine necks | 88,889±19,834 (82,667±18,446) |
| | | Shafts with spines | 96,443±5,006 (89,692±4,656) |
| | | Shafts without spines | 86,178±3,908 (80,146±3,634) |
| | SS | Complete spine heads | 33,283±7,393 (30,953±6,876) |
| | | Incomplete spine heads | 56,528±9,525 (52,571±8,858) |
| | | Spine necks | 33,284±11,330 (30,954±10,537) |
| | | Shafts with spines | 52,584±5,449 (48,903±5,067) |
| | | Shafts without spines | 69,475±10,543 (64,611±9,805) |
| dBA38 | AS | Complete spine heads | 89,839±3,000 (83,551±2,790) |
| | | Incomplete spine heads | 74,253±2,375 (69,055±2,209) |
| | | Spine necks | 49,423±8,486 (45,963±7,892) |
| | | Shafts with spines | 62,475±2,685 (58,101±2,497) |
| | | Shafts without spines | 69,563±3,432 (64,694±3,192) |
| | SS | Complete spine heads | 38,710±5,619 (36,001±5,225) |
| | | Incomplete spine heads | 52,247±9,135 (43,797±7,724) |
| | | Spine necks | 44,662±12,020 (41,536±11,816) |
| | | Shafts with spines | 49,295±3,930 (45,845±3,655) |
| | | Shafts without spines | 60,491±5,332 (56,257±4,959) |

| | | | |
|---|---|---|---|
| **BA21** | AS | Complete spine heads | 133,931±4,024 (124,556±3,743) |
| | | Incomplete spine heads | 130,015±6,739 (120,914±6,267) |
| | | Spine necks | 55,873±7,611 (51,962±7,079) |
| | | Shafts with spines | 83,806±5,074 (76,940±4,719) |
| | | Shafts without spines | 78,436±4,351 (72,945±4,046) |
| | SS | Complete spine heads | 94,452±19,713 (87,840±18,333) |
| | | Incomplete spine heads | 120,534±37,748 (112,097±35,106) |
| | | Spine necks | 15,666±1,970 (14,569±1,832) |
| | | Shafts with spines | 59,744±7,501 (55,562±6,976) |
| | | Shafts without spines | 52,246±7,856 (48,589±7,306) |

**SI Table 6.** SAS area ($nm^2$) of AS and SS established on complete spines heads, incomplete spines heads, spine necks, dendritic shafts with spines and dendritic shafts without spines in layer III of BA24, vBA38, dBA38 and BA21. AS established on complete spine heads were significantly larger than the area of the AS on shafts without spines in BA24 (KW; P<0.05) and vBA38 (KW; P<0.001) — and larger than AS on shafts with spines in dBA38 (KW; P<0.01). Data in parentheses are not corrected for shrinkage. AS: asymmetric synapses; BA: Brodmann's area; d: dorsal; NA: not applicable; SAS: synaptic apposition surface; SE: standard error of the mean; SS: symmetric synapses; v: ventral.

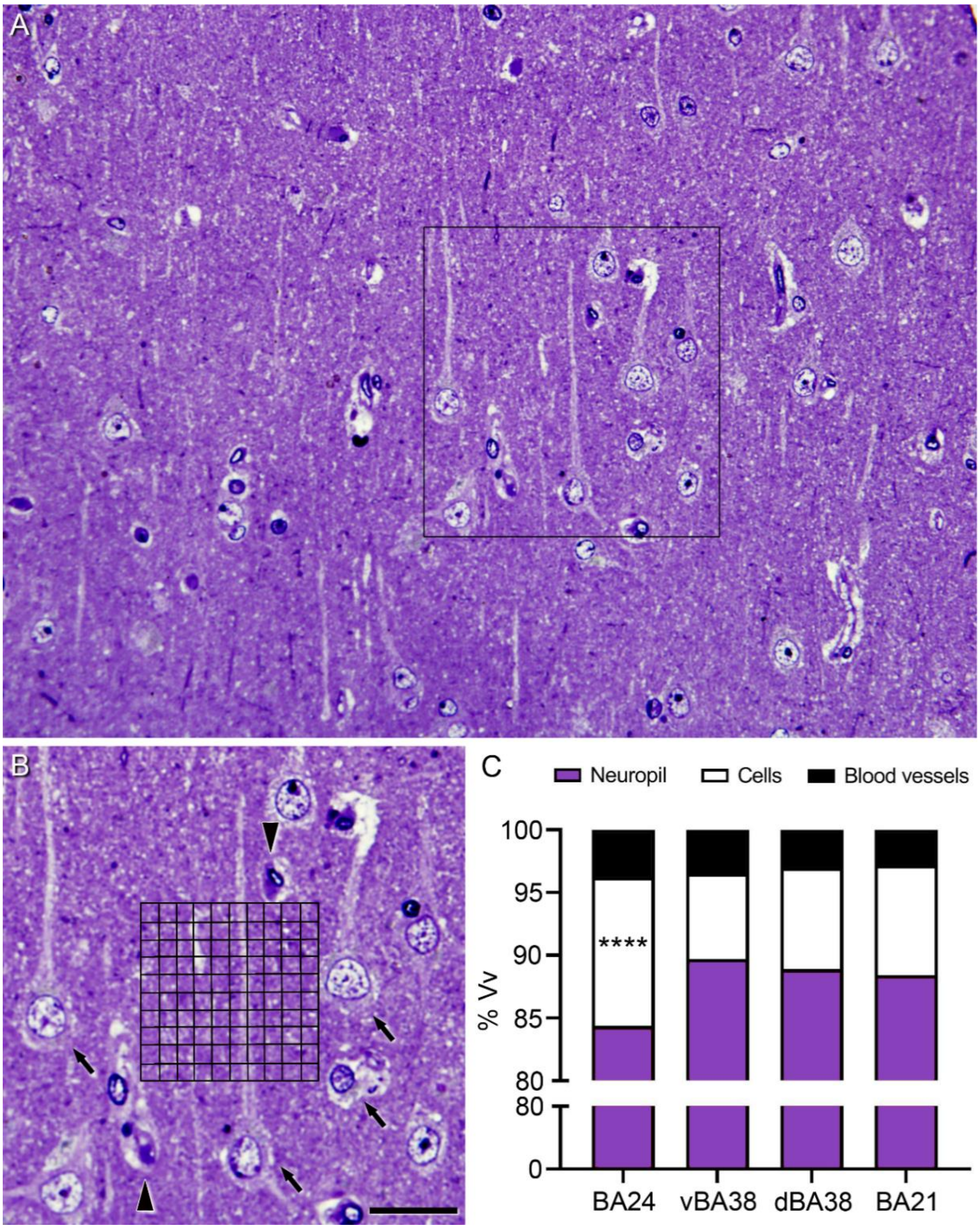


**SI Figure 1.** Stereological estimation of the volume fraction occupied by different cortical elements in BA24, vBA38, dBA38 and BA21. (A) Low-power micrograph of a toluidine blue-stained 1.5 µm-thick semithin section of vBA38. (B) High-power magnification of the boxed area in A showing a 50 µm x 50 µm grid superimposed on the original micrograph, where points hitting the different cortical elements were counted. Some blood vessels (arrowheads) and cell bodies (arrows) are indicated. (C) Plot of the volume fraction occupied by each analyzed cortical element. BA24 shows a higher volume fraction occupied by cells than the other cortical regions ($\chi^2$, $p<0.001$). Scale bar (in B) indicates 40 µm in A, and 25 µm in B. BA: Brodmann's area; d: dorsal; v: ventral.

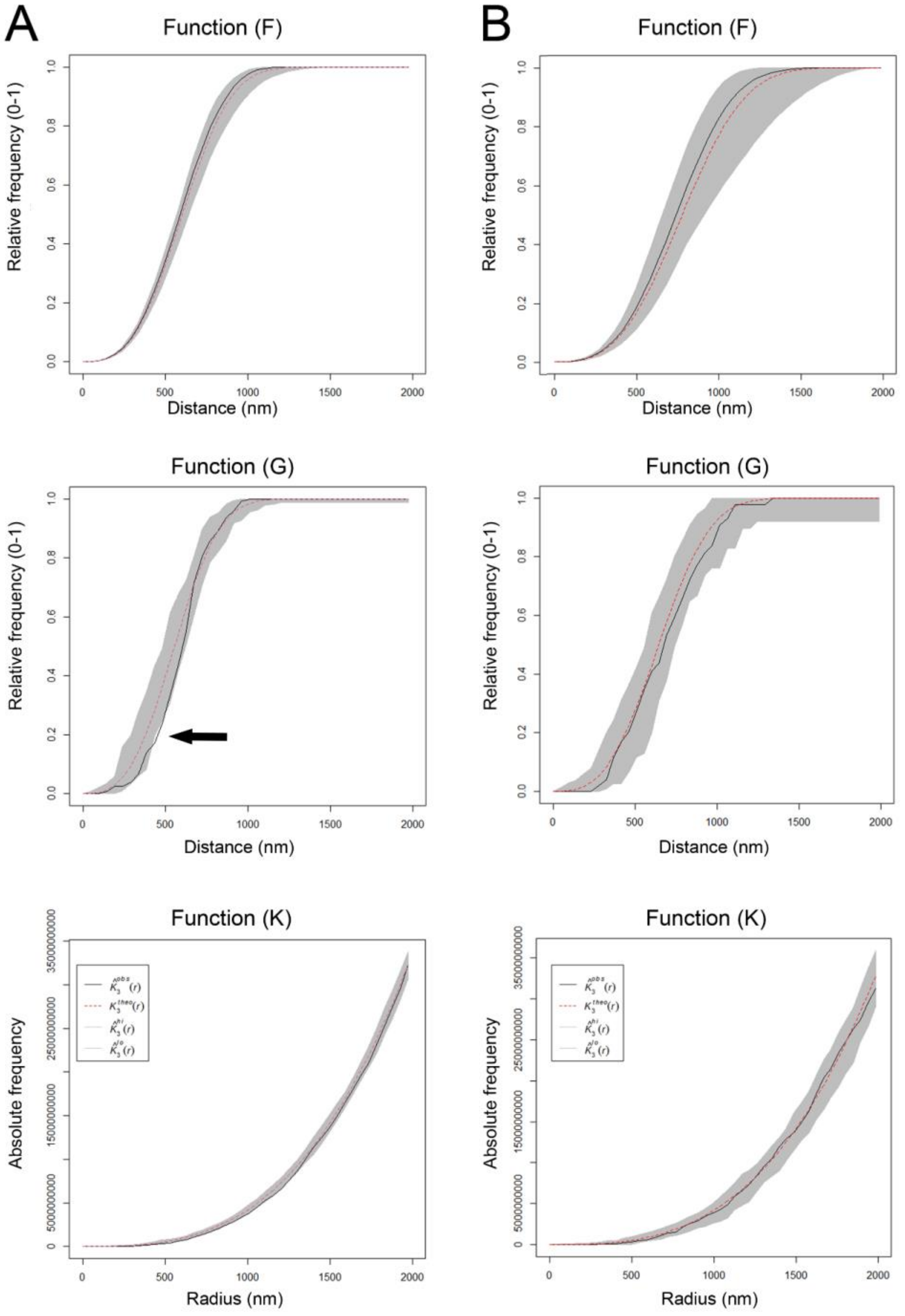


**SI Figure 2.** Analysis of the 3D synaptic spatial distribution in layer III of BA24, vBA38 and dBA38. Red dashed traces correspond to a theoretical homogeneous Poisson process for each function (F, G, K). The black continuous traces correspond to the experimentally observed function in the sample. The shaded areas represent the envelopes of values calculated from a set of 99 simulations. (A) Example of stacks (6 out 27) showing a slight tendency toward a distributed pattern indicated by arrow in G function. (B) Example of stacks (21 out of 27) fitting a Poisson function.

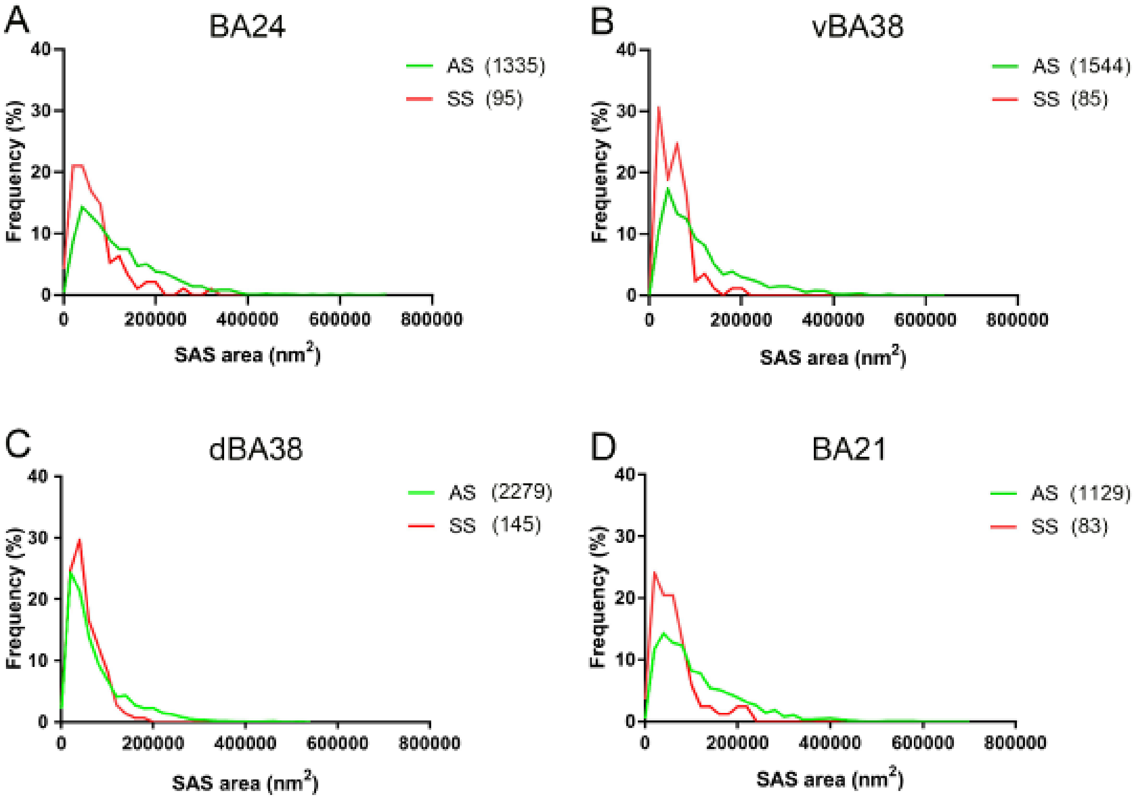


**SI Figure 3.** Frequency distribution of SAS area of AS and SS. (A–D) In all regions, small SS (red) were more frequent than small AS (green) (BA24, KS P<0.0001; vBA38, KS P<0.0001; dBA38, KS P=0.0003; and BA21 KS P<0.0001). AS: asymmetric; BA: Brodmann's area; d: dorsal; SAS: synaptic apposition surface; SS: symmetric; v: ventral.

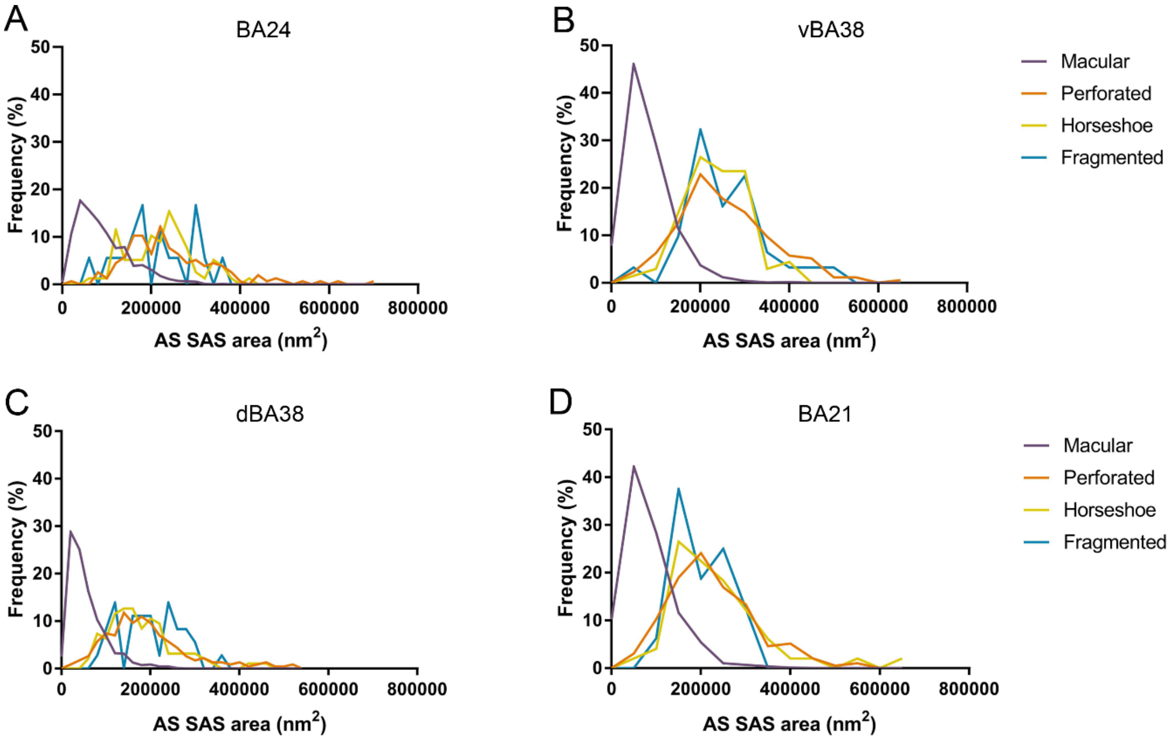


**SI Figure 4.** Frequency histograms of the SAS area of AS per synaptic shape from layer III of BA24 (A), vBA38 (B), dBA38 (C) and BA21 (D). Smaller macular AS were the most frequent shape in all regions (KS, p<0.0001). AS: asymmetric synapses; BA: Brodmann's area; d: dorsal; SAS: synaptic apposition surface; v: ventral.

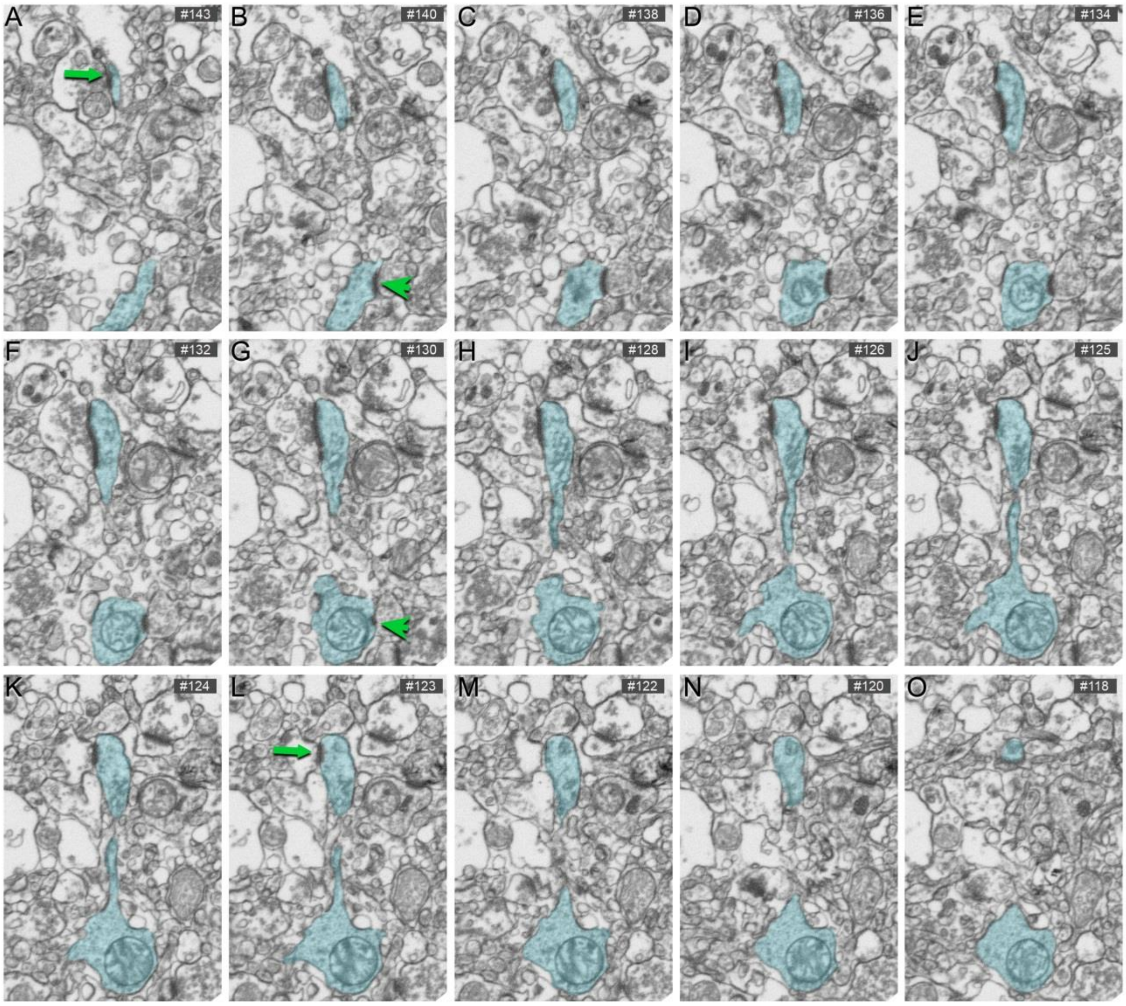


**SI Figure 5.** Serial images obtained by FIB/SEM showing a dendritic segment with a dendritic spine (blue). Asymmetric synapses established on the dendritic spine head (green arrow; A–L) and the dendritic shaft (green arrowhead; B–G) are indicated. Scale bar (in O) = 1 µm in A–O.

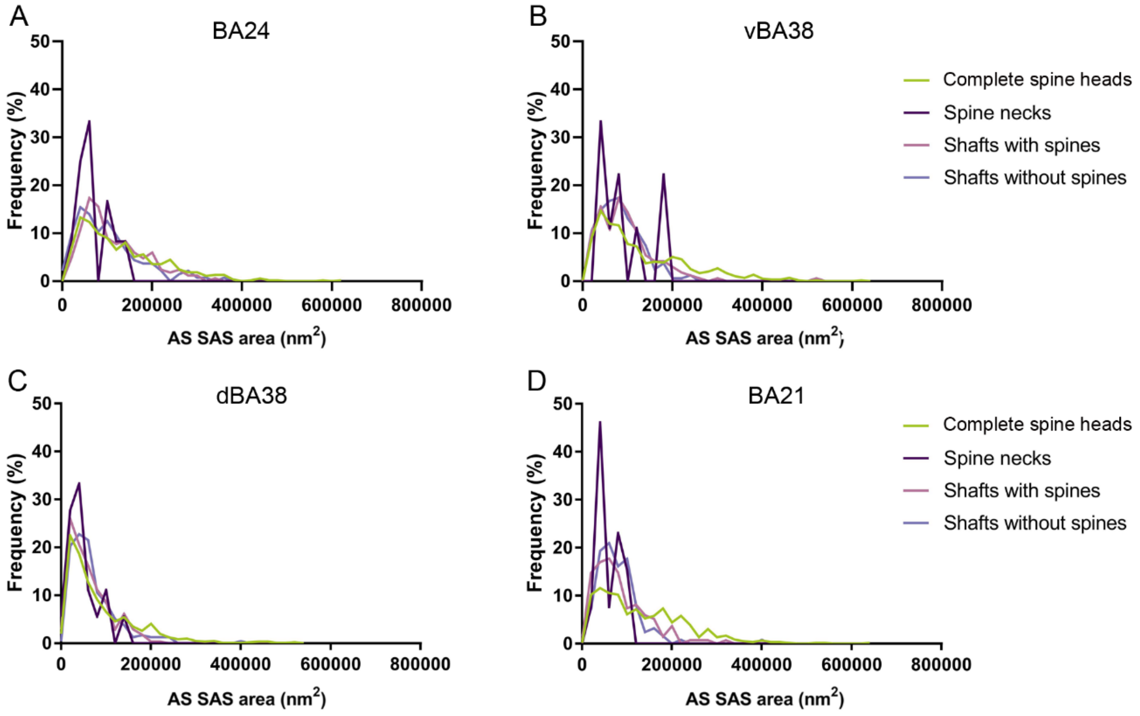


**SI Figure 6**. Frequency histograms of the SAS area of Asymmetric Synapses established on complete spines, spine necks, shafts with spines and shafts without spines from layer III neuropil of BA24 (A), vBA38 (B) dBA38 (C) and BA21 (D) samples. The larger SAS area were significantly more frequent in AS on complete spines than in the case of AS on shafts without spines in BA24 (KS; $P<0.01$), vBA38 (KS; $P<0.0001$), and BA21 (KS; $P<0.0001$) — and more frequent than AS on shafts with spines in dBA38 (KS; $P=0.0002$) and BA21 (KS; $P<0.001$). BA: Brodmann's area; d: dorsal; SAS: synaptic apposition surface; v: ventral.

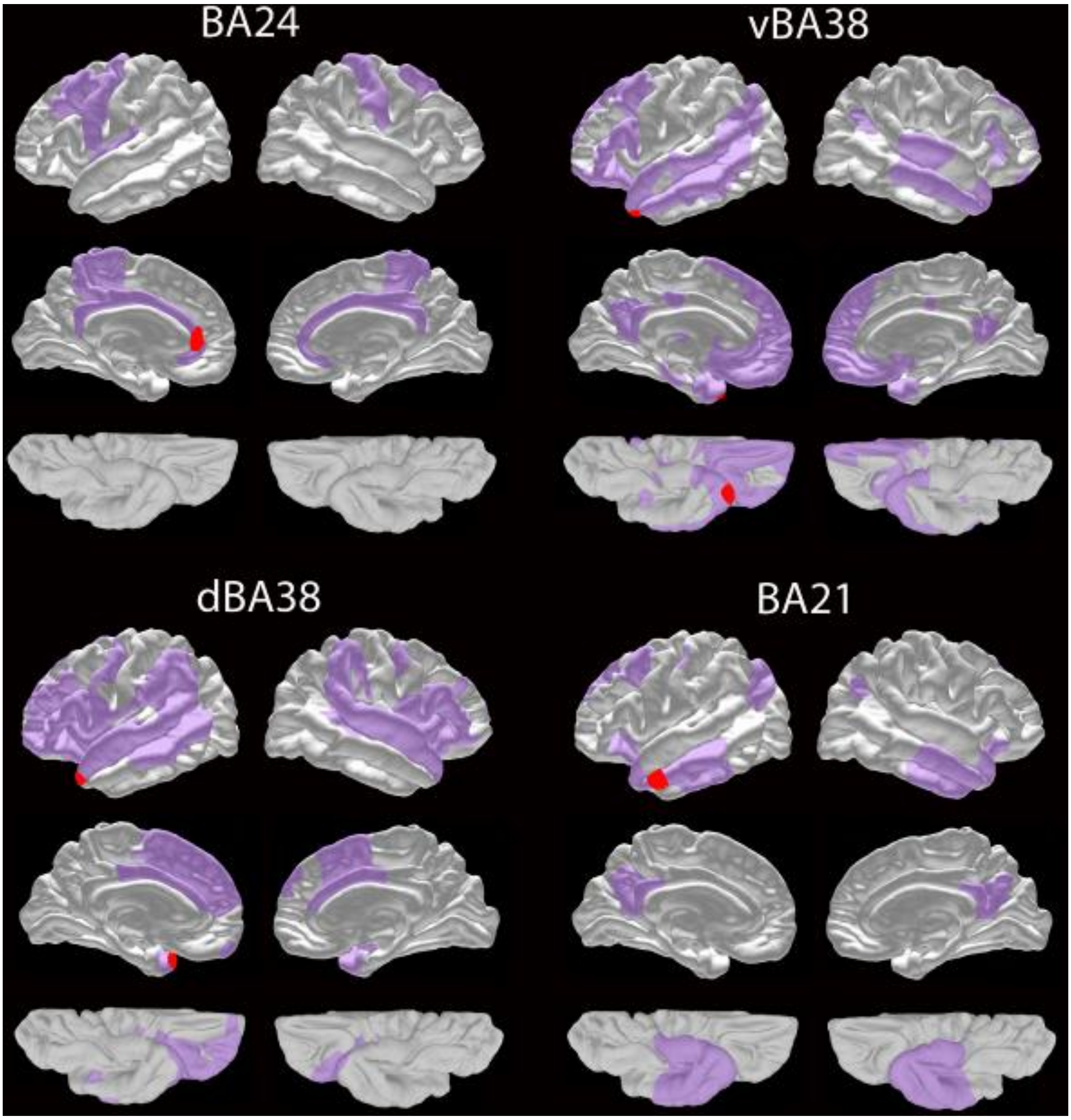


**SI Figure 7.** Schematic representation of cortical connectivity in the analyzed regions. For each region, three views of the left and right hemisphere are shown (lateral, medial and ventral). The location of BA24, vBA38, dBA38 and BA21 are plotted in red, and their connected cortical areas are plotted in purple, indicating the different patterns of intra- and interhemispheric connectivity of the analyzed regions. Data come from the followings functional magnetic resonance imaging studies: BA24 by Jackson et al., (2016); vBA38 and dBA38 by Pascual et al., (2015); and BA21 by Dezachyo et al., (2021). BA: Brodmann's area; d: dorsal; v: ventral. Modified from EBRAINS Interactive Atlas Viewer (https://ebrains.eu/service/human-brain-atlas/#atlas-viewer).

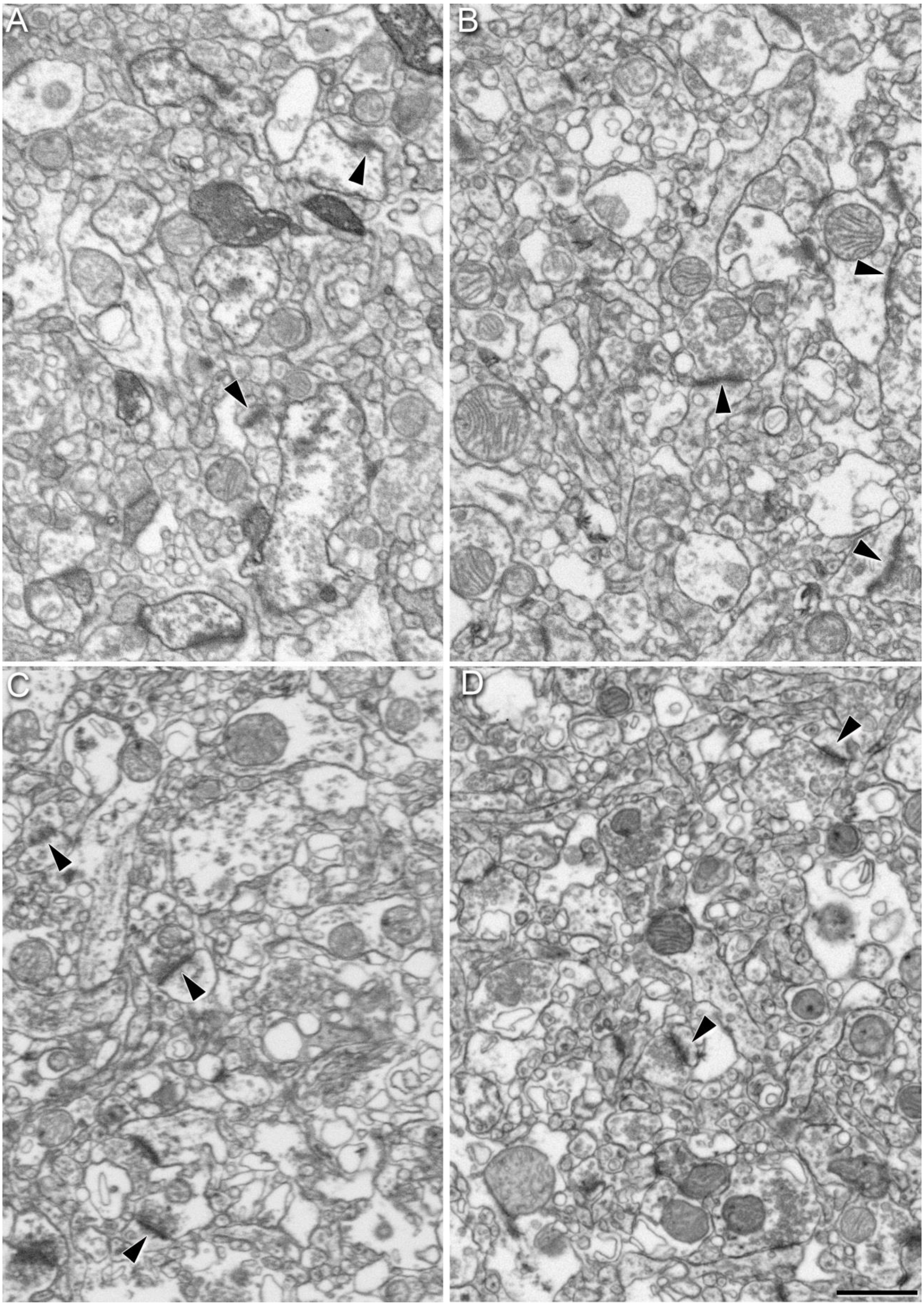


**SI Figure 8.** Images obtained by FIB/SEM showing neuropil of BA24 (A), vBA38 (B), dBA38 (C) and BA21 (D) from an autopsy case (AB3). Some examples of synapses are indicated in all regions (arrowheads). Scale bar (in D) indicates 1 µm in A–D. BA: Brodmann's area; d: dorsal; v: ventral.

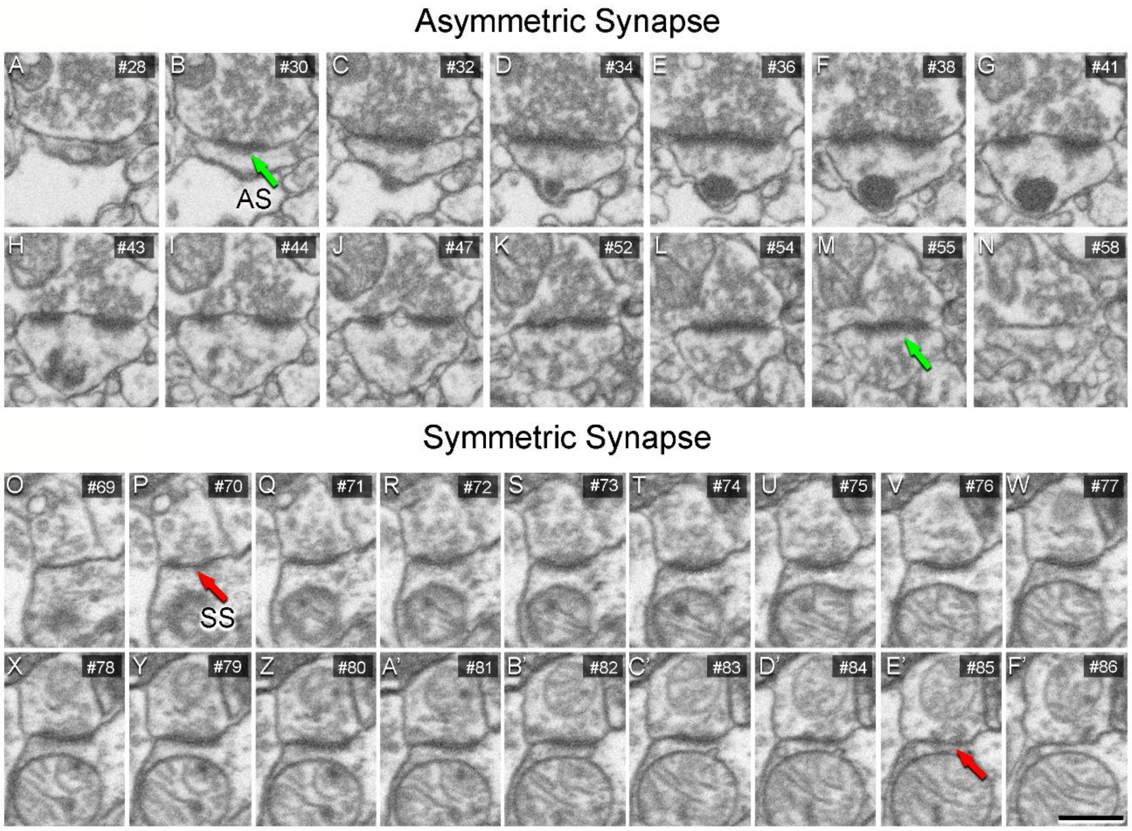


**SI Figure 9.** Sequence of FIB/SEM serial images of an asymmetric synapse (A–N) and a symmetric synapse (O–F'). Numbers on the top right of each panel indicate the number of each section from a stack of serial sections. Synapse classification was based on the examination of full sequences of serial images (see "Three-Dimensional Analysis of Synapses" for further details). Scale bar (in F') indicates 500 nm in A-F'.
